# Crossover designation recruits condensin to reorganize the meiotic chromosome axis

**DOI:** 10.1101/2020.07.16.207068

**Authors:** Victor A. Leon, Tovah E. Markowitz, Soogil Hong, Adhithi R. Raghavan, Jonna Heldrich, Keun Kim, Andreas Hochwagen

## Abstract

Crossover recombination supports meiotic chromosome inheritance and fertility by establishing chiasmata between homologous chromosomes prior to the first meiotic division. In addition to the physical exchange of DNA mediated by meiotic recombination, chiasma formation also involves restructuring of the underlying chromosome axis, possibly to help with chiasma maturation or to resolve chromosomal interlocks. Here, we identify condensin as an important regulator of axis remodeling in *S. cerevisiae*. Condensin is recruited near sites of meiotic crossover designation by pro-crossover factors but is largely dispensable for DNA exchange. Instead, condensin helps to create discontinuities in the meiotic chromosome axis by promoting removal of cohesin. In addition, chromosomes of condensin mutants exhibit unusually common parallel chromatin clouds and experience a chromosomal buildup of the conserved axis remodeler Pch2. Consistent with an important role of axis restructuring at crossover sites, the canonical anaphase-bridge phenotype of condensin mutants is partly rescued by redirecting meiotic DNA repair to sister chromatids instead of homologous chromosomes, suggesting that crossover-associated axis reorganization is important for faithful meiotic chromosome segregation.

## Introduction

Higher-order chromosome architecture allows cells to organize and maneuver large sections of chromatin to modulate gene expression, support DNA metabolism, and enable chromosome segregation ^1,2^. The structural compartmentalization of chromosomes also plays a key role during meiotic recombination ^3–5^. During early meiotic prophase I, chromosomes reorganize into linear arrays of chromatin loops along a protein-rich axis, known as the axial element. This loop-axis arrangement is of central importance for controlling meiotic recombination: it promotes recombination initiation by stimulating the formation of meiotic DNA double-strand breaks (DSBs), ensures the proper targeting of DSBs for homologous repair, and provides a platform for damage surveillance and checkpoint control ^3,6^. In addition, the axial element serves as the foundation for the assembly of the synaptonemal complex (SC), a phase-separated, zipper-like structure that connects the axial elements of paired homologous chromosomes during the later stages of recombination ^4,5^. Crossover intermediates mature within the SC, ultimately leading to the physical exchange of DNA. SC formation, in turn, changes the axial elements, as several axial element proteins become depleted upon SC assembly ^7,8^. Notably, the axial element is also restructured locally at sites of crossover recombination ^9,10^. In the holocentric organism, *C. elegans*, this reorganization determines which chromosome end will attach to the meiotic spindle ^11,12^, whereas in mouse it may help in chiasma maturation ^13^. The reorganization may also facilitate the resolution of chromosomal interlocks that arise as a consequence of meiotic recombination ^9,10^. In several systems, this local reorganization involves a cytologically apparent loss of sister chromatid cohesion ^14–18^. However, the mechanism driving axial element reorganization in the vicinity of crossovers is not understood.

The chromosomal translocase condensin is a central regulator of higher-order chromosome organization in mitosis and meiosis ^19,20^. Condensin belongs to the Structural Maintenance of Chromosomes (SMC) family of protein complexes, which encircle chromatin via a ring-shaped interface formed by two SMC ATPases and a kleisin linker protein. With the help of additional protein subunits, condensin, like other SMC complexes, promotes the ATP-dependent formation of DNA loops ^21,22^. The budding yeast *S. cerevisiae* encodes a single condensin complex that consists of five subunits: the SMC proteins Smc2 and Smc4, the kleisin Brn1, as well as Ycg1 and Ycs4.

Across eukaryotes, condensin is abundantly present on chromosomes at the time of meiotic recombination ^23–26^, yet its precise function remains poorly understood. In *S. cerevisiae* and *S. pombe*, condensin suppresses recombination of repetitive DNA ^27,28^ while in *C. elegans*, it controls the distribution of meiotic DSBs ^29–31^. However, condensin is not required for normal DSB repair kinetics ^23^. In late prophase I, condensin promotes the reduction of cohesin, another SMC complex, from meiotic chromosomes axes ^32^, and contributes to SC disassembly and chromosome individualization ^29,33–35^. Similar to the characteristic chromosome bridging phenotype in mitosis, condensin mutants also exhibit chromosome bridges during meiosis, partly due to an inability to efficiently remove cohesin ^23,32^. Intriguingly, the formation of meiotic chromosome bridges in condensin mutants is also partially dependent on meiotic DSB formation ^23^, but the basis for this DSB-dependence has remained elusive.

Here, we further analyzed the function of condensin during meiotic prophase I in *S. cerevisiae*. We show that crossover designation triggers condensin enrichment near sites of crossover repair. Loss of condensin function led to more continuous axial elements, a higher level of cohesin at crossover-designated sites, and an accumulation of the axis remodeler Pch2. Importantly, targeting recombination events to sister chromatids rather than homologous chromosomes was sufficient to reduce meiotic anaphase bridging. These findings suggest that condensin-mediated reorganization of the axial element at sites of interhomolog recombination is important for ensuring faithful meiotic chromosome segregation.

## Results

### Condensin binds to the axial element and DSB hotspots in response to recombination initiation

To define the mechanism of condensin accumulation on chromosomes during meiotic prophase I, we induced cultures to undergo a synchronous meiotic time course and analyzed the distribution of the condensin SMC subunit Smc4, functionally fused to GFP (**Figure S1A**) on chromosome spreads. In early prophase (2 hours), Smc4-GFP showed some foci on non-nucleolar chromatin but was primarily enriched in the nucleolus as indicated by a cluster of foci near the nucleolar marker Nop1 (**Figure 1A-C**). As cells progressed into meiotic prophase (3-4 hours) and initiated SC formation, the nucleolar enrichment of Smc4-GFP weakened, and condensin formed numerous foci along chromosome axes that localized on or adjacent to the SC protein Zip1 (**Figure 1A-B, Figure S1B**), consistent with prior analysis ^23^.

**Figure 1.**
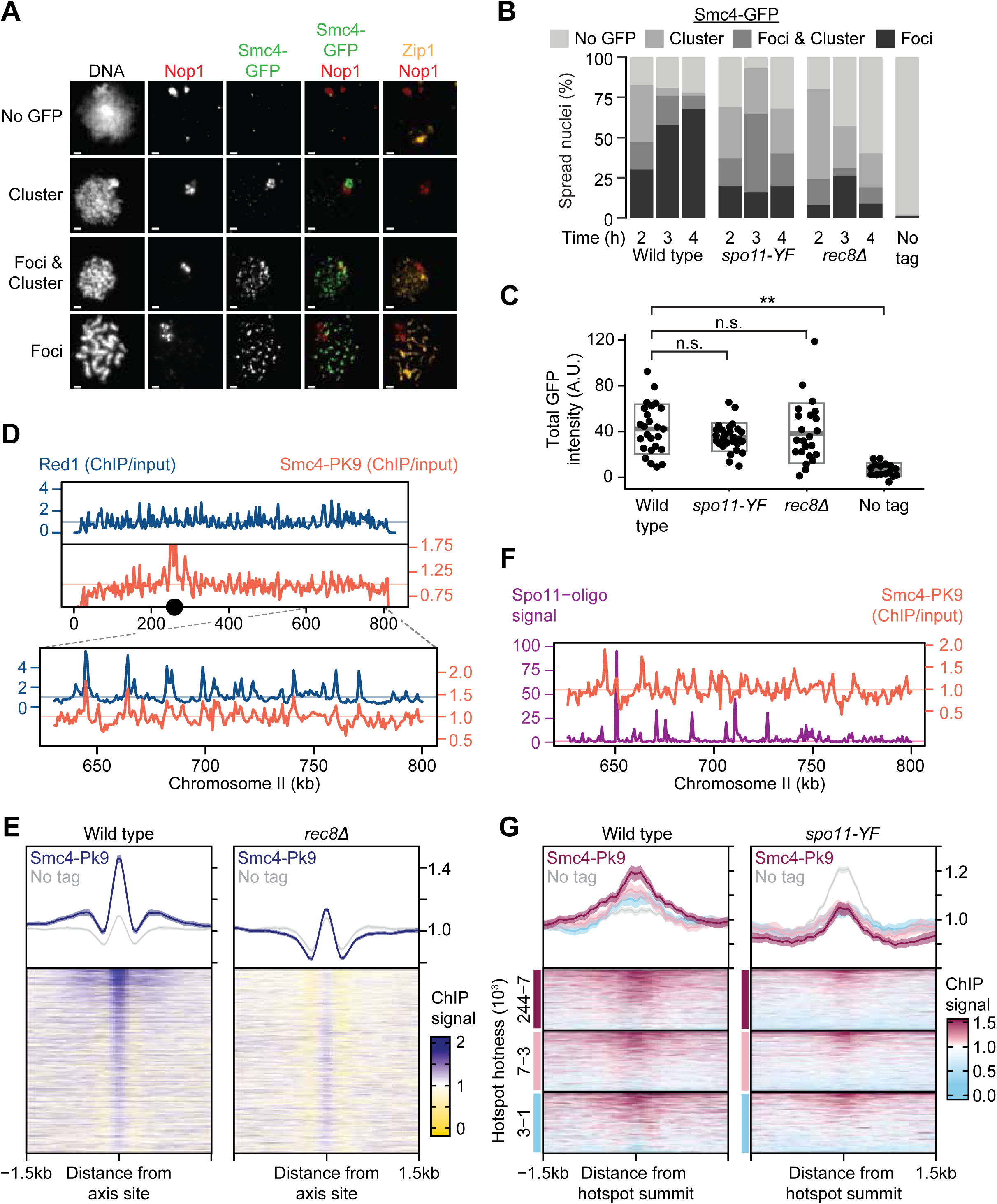
Condensin distribution on prophase chromosomes depends on *SPO11* and *REC8*. (**A**) Representative images of Nop1 (red), Smc4-GFP (green), and Zip1 (orange) binding patterns on surface-spread nuclei, categorized as “No GFP”, “Foci”, “Foci & Cluster”, or “Cluster” of GFP patterns. Bar = 1 µm. (**B**) Incidence of GFP patterns shown in (A) in wild-type [H9443], *rec8*Δ [H9442], *spo11- YF* [H9441], or in the absence of a GFP tag (“no tag”) [H7797], n=100. (**C**) Quantifications of GFP levels on nuclei from wild-type, *rec8*Δ, *spo11-YF*, and untagged control samples 3h after meiotic induction. Only nuclei with visible Zip1 staining were analyzed, n=25, 24, 27, 16 respectively. Total GFP intensity for each nucleus was normalized to background. Bars indicate mean and standard deviation (S.D). Wilcox test (alternative hypothesis: greater). (**D**) Red1 (blue) [H119/6408] and Smc4-PK9 (pink) [H6408] binding patterns along chromosome II and a representative region (inset) in wild-type cells. Horizontal blue and pink lines represent genome averages. (**E**) Heatmap of Smc4-PK9 enrichment around axis sites in wild type [H6408] and *rec8*Δ mutants [H7660]. Axis sites were defined as Red1 summits in wild-type cells with -log10 q-value greater than or equal to 20, as determined by MACS2. ChIP-seq values were averaged over 50-bp windows around axis sites across a 3-kb window. Rows are sorted by amount of Red1 binding at these sites. Heatmap colors represent enrichment (blue) or depletion (gold). Mean and 95% confidence intervals are shown in line graphs directly above heatmap. Gray line indicates mean and standard error of the mean (S.E.M.) from the no-tag controls [H7797, H8428] and provides a measure of the non-specific enrichment by the PK9 antibody. (**F**) Spo11-oligo signal (purple) ^46^ and Smc4-PK9 (pink) [H6408] binding pattern in wild-type cells in the same representative region on chromosome II shown in (D). Horizontal lines represent genome averages. (**G**) Heatmap of Smc4-PK9 enrichment around DSB hotspots in wild-type cells [H6408] and *spo11-YF* mutants [H8630]. Hotspots were placed into three quantiles grouped by strength (from hottest to weakest: maroon, pink, light blue). ChIP-seq values were averaged over 50-bp windows around the midpoints of hotspots across a 3-kb window. Rows in each quantile were sorted by Smc4-PK9 enrichment with pink indicating high enrichment and light blue indicating depletion. Mean and 95% confidence intervals are shown in the line graph directly above the heatmap. Gray lines indicate mean and 95% confidence intervals of the no-tag controls [H7797, H8643]. The signal seen at hotspots in no-tag controls is because hotspots coincide with nucleosome-free regions, which fragment more frequently during sonication and thus create non-specific background signals. See also **Figure S1**.

To identify chromosomal sequences enriched for condensin, we analyzed Smc4 functionally tagged with PK9 ^36,37^ (**Figure S1A**) by chromatin immunoprecipitation and sequencing (ChIP-seq). Synchronous cell populations were collected 3 hours after meiotic induction, corresponding to mid-prophase I. Initial experiments using a standard ChIP-seq protocol failed to recover specific peaks of condensin enrichment when compared to a no-tag control (**Figure S1C**). The only exception was a specific peak in the ribosomal DNA (rDNA; not shown). This recovery problem was meiosis-specific ^37^ and appeared to be linked to poor fragmentation of condensin-associated chromatin: proteolytic elimination of proteins after the immunoprecipitation step and re-sonication of the pure DNA fragments prior to library preparation ^38^ led to the recovery of numerous specific peaks of Smc4-PK9 enrichment. These peaks were absent in a similarly treated control strain lacking the PK9 tag (**Figure S1C**).

Binding profiles of Smc4-PK9 from re-sonicated meiotic samples exhibited several features previously observed for condensin in vegetative cells, including enrichment at tRNA genes, around centromeres, and within the rDNA ^36,39^ (**Figure S1D-F**). We also detected condensin accumulation between *CEN12* and the rDNA, consistent with prior observations in mitotic cells ^37^ (**Figure S1G**). In vegetative cells, condensin enrichment at pericentromeres and next to the rDNA is only observed during mitosis ^36,40–42^. The presence of similar enrichment in meiotic cells thus likely reflects the high degree of chromosomal compaction that characterizes meiotic prophase I ^23^. Importantly, we identified meiosis-specific condensin enrichment patterns, most prominently at sites of axial element attachment and at DSB hotspots (**Figure 1D-G**). Axis attachment sites are defined by strong local enrichment of the meiosis-specific cohesin subunit Rec8 and the axial element factor Red1 ^43–45^. Although Smc4-PK9 peaks visibly outnumbered the peaks of Rec8 and Red1, the large majority of Rec8 and Red1 peaks overlapped with Smc4-PK9 peaks (**Figure 1D-E, Figure S1E)**. Many of the remaining meiosis-specific peaks localized to hotspots of DSB formation where condensin enrichment correlated broadly with hotspot activity (**Figure 1F-G**) ^46^.

To directly test whether condensin binding is coordinated with meiotic recombination, we analyzed *spo11-Y135F* (*spo11-YF*) mutants, which lack catalytically active Spo11 nuclease and are unable to initiate meiotic DSB formation ^47,48^. Although Smc4-GFP binding remained detectable on chromosomes in these mutants, the level of chromosomal association did not change over time and binding remained strong in the nucleolus (**Figure 1B**). In addition, Smc4-PK9 binding at recombination hotspots became nearly undetectable in *spo11-YF* mutants (**Figure 1F**), indicating that condensin enrichment on meiotic chromosomes is largely dependent on the initiation of meiotic recombination.

Recruitment of Smc4 to meiotic chromosomes was also strongly reduced in *rec8*Δ mutants (**Figure 1B, E**). Because loss of *REC8* causes reduced DSB levels in some regions of the genome ^43,49^, we considered the possibility that the reduced Smc4 binding was secondary to reduced DSB formation. However, analysis of individual chromosomes revealed that, in contrast to the region-specific defects in DSB formation, Smc4 binding was uniformly impacted across chromosomes in *rec8*Δ mutants (**Figure S1E**). This observation suggests that condensin recruitment requires the presence of Rec8-cohesin independently of its role in promoting DSB formation. Notably, total signal intensity of chromosome spreads with detectable Smc4-GFP signal was not significantly different between wild type, *spo11-YF* mutants, and *rec8*Δ mutants (**Figure 1C, S2A**), indicating that the changes in condensin patterns were primarily the result of condensin redistribution from the nucleolus to chromosomes rather than altered protein expression or stability. We conclude that meiotic DSB formation directs condensin enrichment on chromosome arms and that both DSB hotspots and Rec8-dependent chromosomal processes provide binding sites for condensin upon DSB formation in meiotic prophase.

### DSB-dependent condensin recruitment requires crossover designation but not synapsis

In *S. cerevisiae*, DSB formation triggers a series of changes along meiotic chromosomes that culminate in the assembly of the SC ^50,51^. Previous work had indicated that condensin remains bound to meiotic chromosomes in mutants lacking the SC transverse filament protein Zip1 ^23^. However, as condensin redistribution coincided with the appearance of short stretches of SC (**Figure S1B**), we decided to revisit the possibility that condensin redistribution is linked to SC assembly. Analysis of *zip1*Δ mutants revealed Smc4-GFP signal along Red1-decorated axial elements (**Figure 2A-B**). However, unlike in wild-type cells, Smc4 also remained abundantly enriched in the nucleolus. In fact, the relative amounts of nucleolar and non-nucleolar staining were not significantly different between *zip1*Δ and *spo11-YF* mutants (**Figure 2A-B**). A similar pattern was observed in *zip2*Δ mutants (**Figure S2B-D**), which fail to load Zip1 onto chromosome arms ^52^. In contrast, disruption of the central DNA-damage sensor kinase Mec1, which mediates many DSB-dependent signals in meiotic prophase, did not affect Smc4 redistribution (**Figure S2E**). Total protein levels of Smc4-GFP and another condensin subunit Ycs4-13myc remained unchanged in *zip1*Δ and *zip2*Δ mutants (**Figure S2A-B**) ruling out differences in expression or stability. Together, these data indicate that DSB-dependent redistribution of condensin from the nucleolus to chromosomes depends on Zip1.

**Figure 2.**
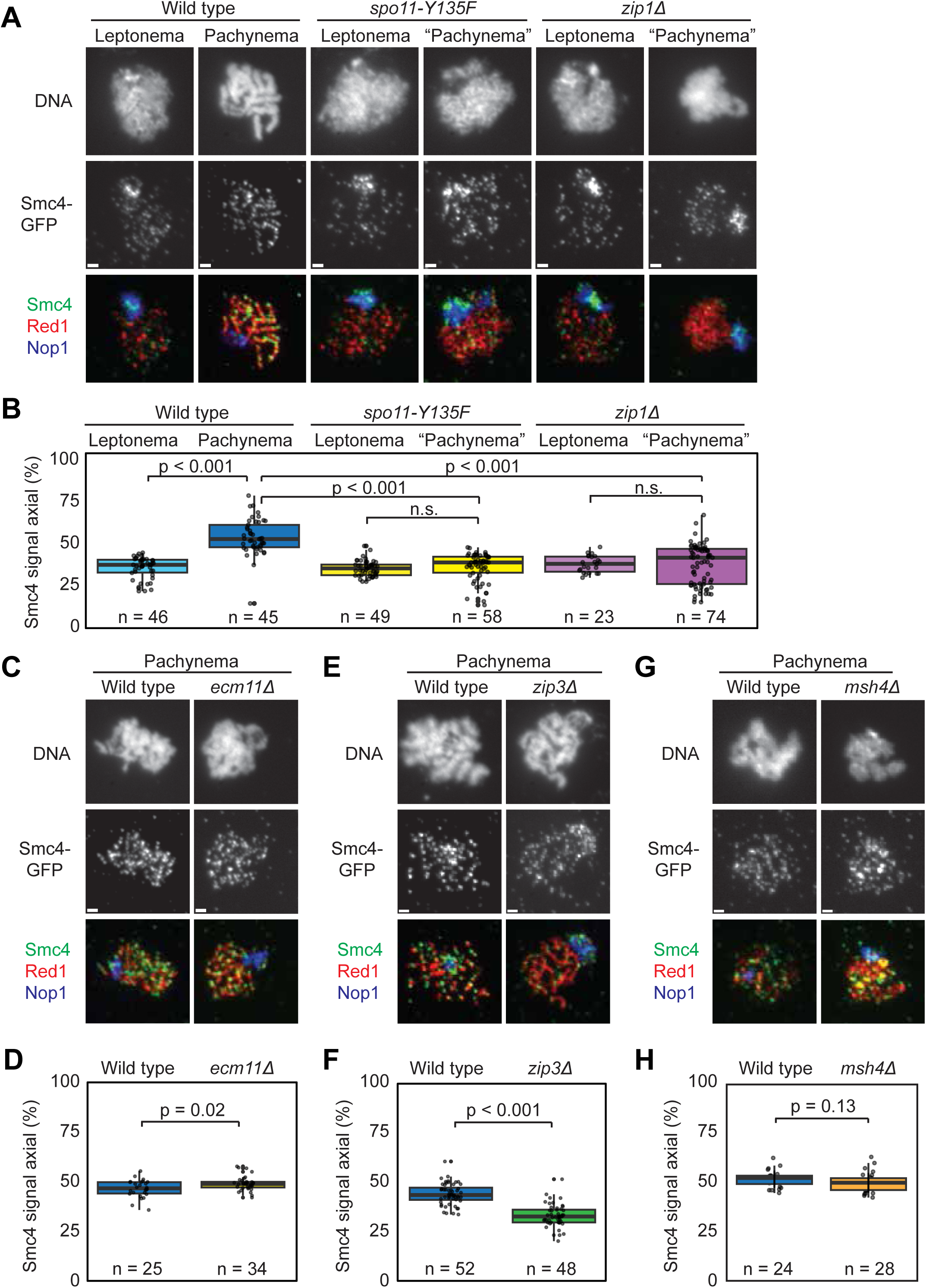
Increased condensin binding to meiotic chromosomes requires crossover designation. (**A**) Representative images of Smc4-GFP (green) distribution on surface-spread nuclei from wild-type [H11471], *spo11-YF* [H9441], and *zip1*Δ mutants [H11499]. Chromosomes were also stained for DNA (DAPI), Red1 (red), and the nucleolar marker Nop1 (blue). Nuclei were staged based on Red1 staining. Punctate Red1 staining was classified as leptonema and individualized Red1 axes were classified as pachynema (note that pachynema is defined by complete synapsis, which does not occur in *spo11-YF* and *zip1*Δ mutants; however, individualized Red1 axes are typically associated with pachynema in wild-type cells). Bar = 1 µm. (**B**) Smc4-GFP signal overlapping with Nop1 (nucleolar) or not overlapping with Nop1 (axial) was quantified. The percent of axial signal out of total signal is shown. Statistical significance was determined using Welch two-sample *t*-tests with Benjamini-Hochberg correction. (**C-G**) Representative images of Smc4-GFP on spread wild-type [H11471] and mutant chromosomes as well as quantification of relative Smc4 distribution in pachynema. (**C-D**) *ecm11*Δ [H12818], (**E-F**) *zip3*Δ [H12925], **(G-H**) *msh4*Δ [H13168]. See also **Figure S2**.

Zip1 is a multifunctional protein that, in addition to supporting chromosome synapsis, also functions as part of the ZMM proteins to designate sites of crossover repair and protect nascent crossover repair intermediates from dissolution ^53–56^. To distinguish whether condensin recruitment relies on synapsis or crossover designation, we analyzed mutants that disrupt these processes individually. Smc4-GFP recruitment was unaffected in *ecm11*Δ mutants (**Figure 2C-D**), which recombine normally but are unable to synapse chromosomes ^57^. By contrast, condensin redistribution was strongly defective in *zip3*Δ and *zip4*Δ mutants, two other *zmm* mutants that are competent for partial SC formation (**Figure 2E-F, Figure S2C-D**) ^58,59^. Thus, condensin recruitment to meiotic chromosome arms requires ZMM-dependent crossover designation but not synapsis.

The ZMM group of proteins includes not only factors required for crossover designation and synapsis initiation (Zip1-4) but also subunits of the MutSγ complex (Msh4 and Msh5), which is thought to stabilize nascent crossover intermediates ^60^. Intriguingly, Smc4-GFP localized to chromosome arms in *msh4*Δ and *msh5*Δ mutants but formed abnormal aggregates on chromosomes (**Figure 2G-H, Figure S2C-D**). These observations indicate that, unlike Zip1-4, MutSγ is not required for condensin recruitment *per se*, but plays a role in controlling condensin focus formation.

To determine whether condensin localizes to sites of crossover designation, we compared the distribution of Smc4-GFP with Zip3 and Msh4, both tagged with 13myc. These proteins are well-established markers of crossover designation and repair ^56,58,61^. We detected a large fraction of condensin foci at a consistent distance from Zip3 and Msh4 foci (**Figure 3A-F**). In contrast, no such association was observed between Smc4-GFP and the axis remodeler Pch2, which also forms numerous foci along synapsed chromosomes (**Figure S2F**) ^62,63^. These results support the conclusion that ZMM factors are required for the recruitment and proper accumulation of condensin near sites of crossover designation.

**Figure 3.**
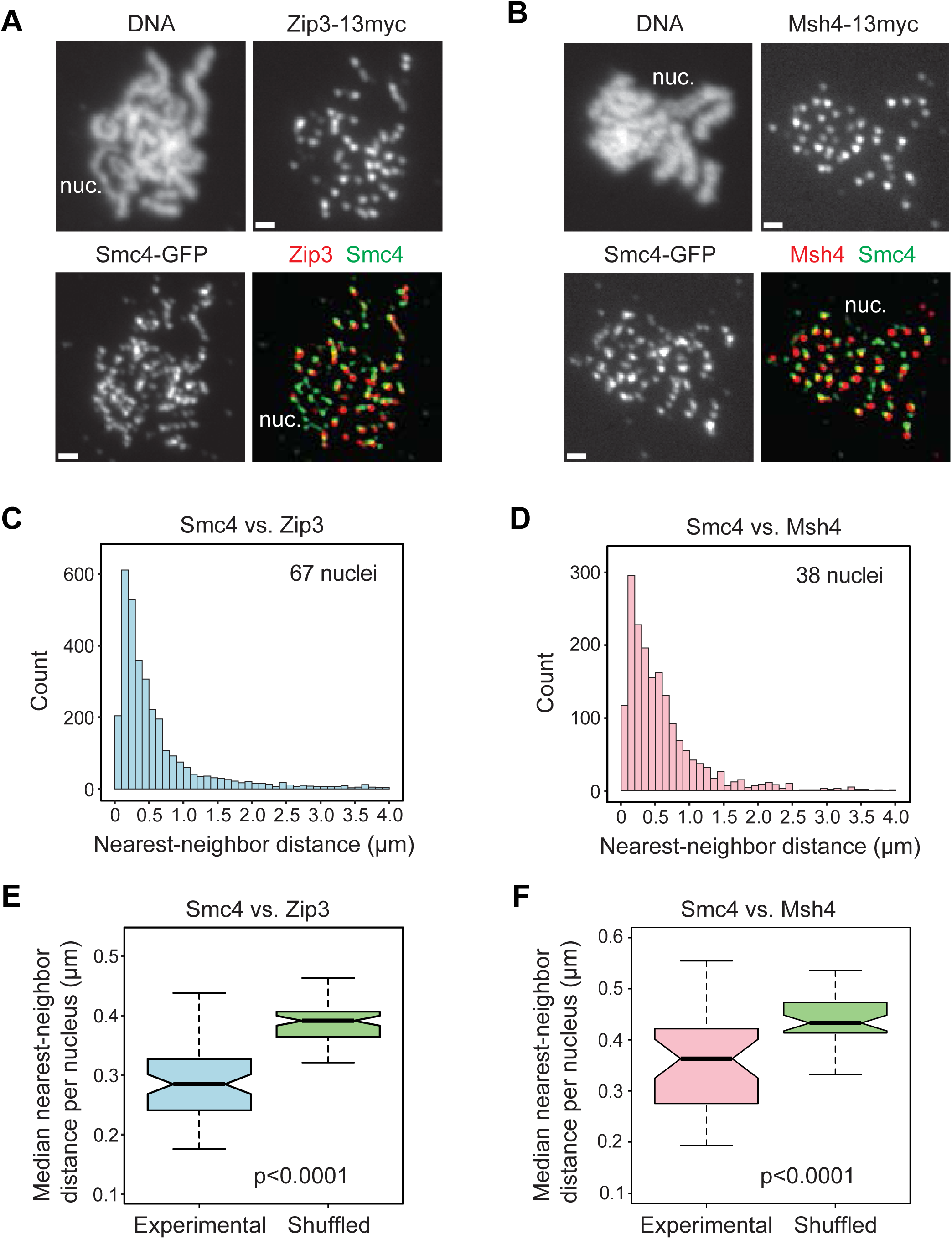
Condensin is enriched near sites of crossover designation. Representative images showing the relative distribution of Smc4-GFP (green) compared to (**A**) Zip3-13myc [H12926] or (**B**) Msh4-13myc [H11735] on surface-spread pachytene nuclei. DNA was stained with DAPI. nuc. = nucleolus, as apparent from the unsynapsed chromatin. (**C-D**) Distance analysis (DiAna) of Smc4-GFP and Zip3-13myc (C) and Msh4-13myc (D), respectively. The distances between nearest-neighbor foci were measured using focus centers. Bar graphs show the combined nearest-neighbor focus distances collected from the indicated number of nuclei. (**E-F**) Distribution of median focus distances per nucleus for Smc4/Zip3 focus pairs (E) and Smc4/Msh4 focus pairs (F). Input data is the same as for (C) and (D) and compared to matched randomized controls (see Methods). Wilcoxon rank sum exact test.

### Condensin is not a ZMM factor

The ZMM-dependent recruitment of condensin raised the possibility that condensin itself may function as a ZMM factor. Classical *zmm* mutants share several diagnostic phenotypes, including a strong prophase arrest in meiotic prophase at high temperatures (33°C), reduced crossover formation, and increased non-crossover repair events ^54^. To test whether condensin acts as a ZMM factor, we analyzed meiotic progression in several temperature-sensitive condensin mutants. The mutants were allowed to enter the meiotic program at 25℃ before shifting them to 34℃, the maximal temperature for normal meiotic progression in our strain background, around the time of early premeiotic S phase (1h time point).

*ycg1-2* and *smc2-8* mutants completed meiosis efficiently at 25℃ but failed to form spores when shifted to 34℃ (**Figure S3A-B**). *ycs4-2* cells already showed sporulation defects at room temperature, indicating that 25℃ is partially restrictive for this mutant. By contrast, *smc4-1* mutants only showed a partial sporulation defect at 34℃, implying that *smc4-1* has a higher restrictive temperature (**Figure S3A-B**). In line with the sporulation defect, condensin mutants at 34℃ failed to properly separate DNA masses during the meiotic divisions (see e.g. **Figure S4E**), corroborating previous studies ^23,32^. Importantly, however, condensin mutants formed the meiosis I spindle with wild-type kinetics at 34℃ (**Figure S3B**). Thus, unlike *zmm* mutants, inactivation of condensin does not lead to a strong prophase arrest. The exception was *ycs4-2*, which showed asynchronous meiotic progression at both temperatures. Given the severity of this allele already at 25℃, these delays may be due to defects originating in the preceding mitotic divisions.

To further distinguish condensin from ZMM factors, we analyzed meiotic DSB formation and repair in condensin mutants at the restrictive temperature. Pulsed-field gel analysis revealed largely normal levels and distribution of meiotic DSBs in *ycg1-2* mutants along chromosome III (**Figure S3C**), in line with previous analyses of a major DSB hotspot ^23^. At the model *HIS4LEU2* locus, DSB formation appeared slightly delayed in *ycg1-2* and *smc2-8* mutants (**Figure 4A-B**). However, crossover repair progression, including single-end invasion, double Holliday junction formation, and crossover resolution occurred at wild-type levels in both mutants (**Figure 4A, 4C-D, Figure S4A-D**). Non-crossovers formation was similarly unaffected (**Figure 4E-F**). Together, these data indicate that although condensin requires ZMM factors for its recruitment near crossover sites, condensin itself is not a ZMM factor.

**Figure 4.**
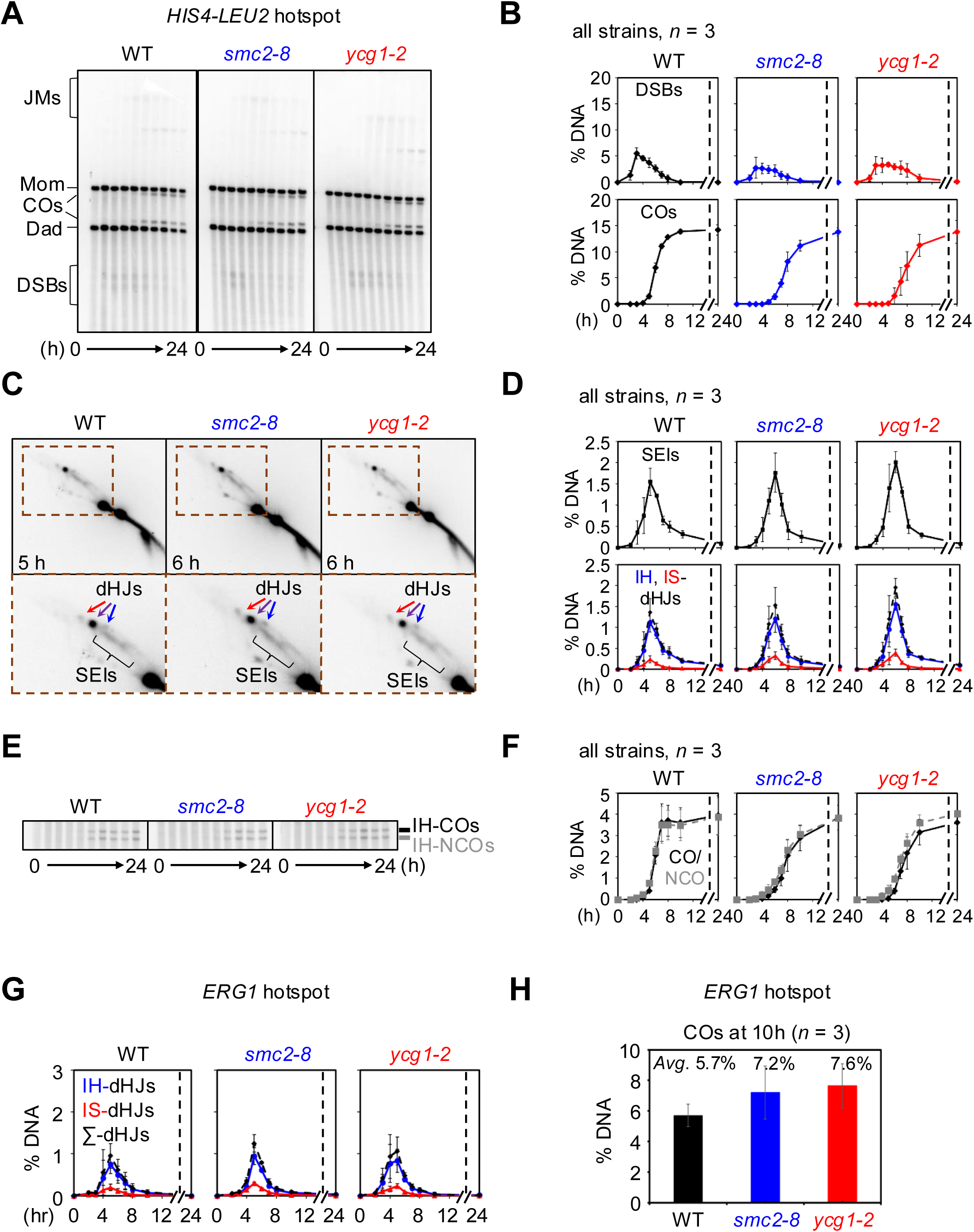
Largely normal joint-molecule and crossover levels in condensin mutants. Analysis of meiotic recombination at the engineered *HIS4-LEU2* hotspot (**A-F**) and the natural *ERG1* hotspot (**G-H**) in a synchronous meiotic time course. Cultures were shifted to 34°C after 2.5 hours (see Methods and **Figure S4** for details). (**A**) Representative 1D gel images of wild-type [KKY4278], *smc2-8* [KKY6547], and *ycg1-2* mutant strains [KKY6544]. Double-strand break, DSB; joint molecules, JMs; crossovers, COs. (**B**) Quantification of DSB and CO species observed in (A). (**C**) Representative 2D gel images of wild-type, *smc2-8*, and *ycg1-2* mutant strains. (**D**) Quantification of single-end invasion (SEI), interhomolog double Holliday junction (IH-dHJ), and intersister double Holliday junction (IS-dHJ) species from (C). (**E**) Representative 1D gel images showing interhomolog crossovers (IH-CO) and interhomolog non-crossovers (IH-NCO) in wild-type, *smc2-8*, and *ycg1-2* mutant strains. (**F**) Quantification of IH-COs and IH-NCOs from (E). (**G**) Quantification of IH-dHJ and IS-dHJ species at *ERG1* in wild-type, *smc2-8*, and *ycg1-2* mutant strains. (**H**) Quantification of CO levels at *ERG1* in wild-type, *smc2-8*, and *ycg1-2* mutant strains 10 hours after meiotic induction. See also **Figures S3-S4**.

Because DSB levels at *HIS4LEU2* are extraordinarily high, we also analyzed crossover formation at the natural *ERG1* hotspot ^46^. At this hotspot, joint molecule formation was mildly elevated in condensin mutants, leading to an approximately 15% increase in crossover formation (**Figure 4G-H, Figure S4F-H**). This increase in recombination activity may be linked to the mild synapsis defects of condensin mutants ^23^ (see below), as similar increases in crossover formation are also observed in mutants lacking the SC central element components Ecm11 and Gmc2 ^64^.

### Condensin is required for normal meiotic axis morphogenesis

Our data suggest that condensin performs functions near crossover-designated sites that are not related to DNA repair. To further investigate this possibility, we analyzed the morphology of crossover-designated sites in condensin mutants using Zip3-GFP as a marker. These analyses included a meiosis-specific depletion allele of the condensin subunit Brn1 (*brn1-mn*), which allows efficient Brn1 depletion after shift to sporulation medium ^65^. While the number of Zip3 foci remained largely unaltered in condensin mutants (**Figure 5A, Figure S5**), consistent with the overall normal levels of crossover formation, the size and intensity of Zip3 foci were significantly increased (**Figure 5A-C**), indicating that condensin shapes the architecture of crossover-designated sites.

**Figure 5.**
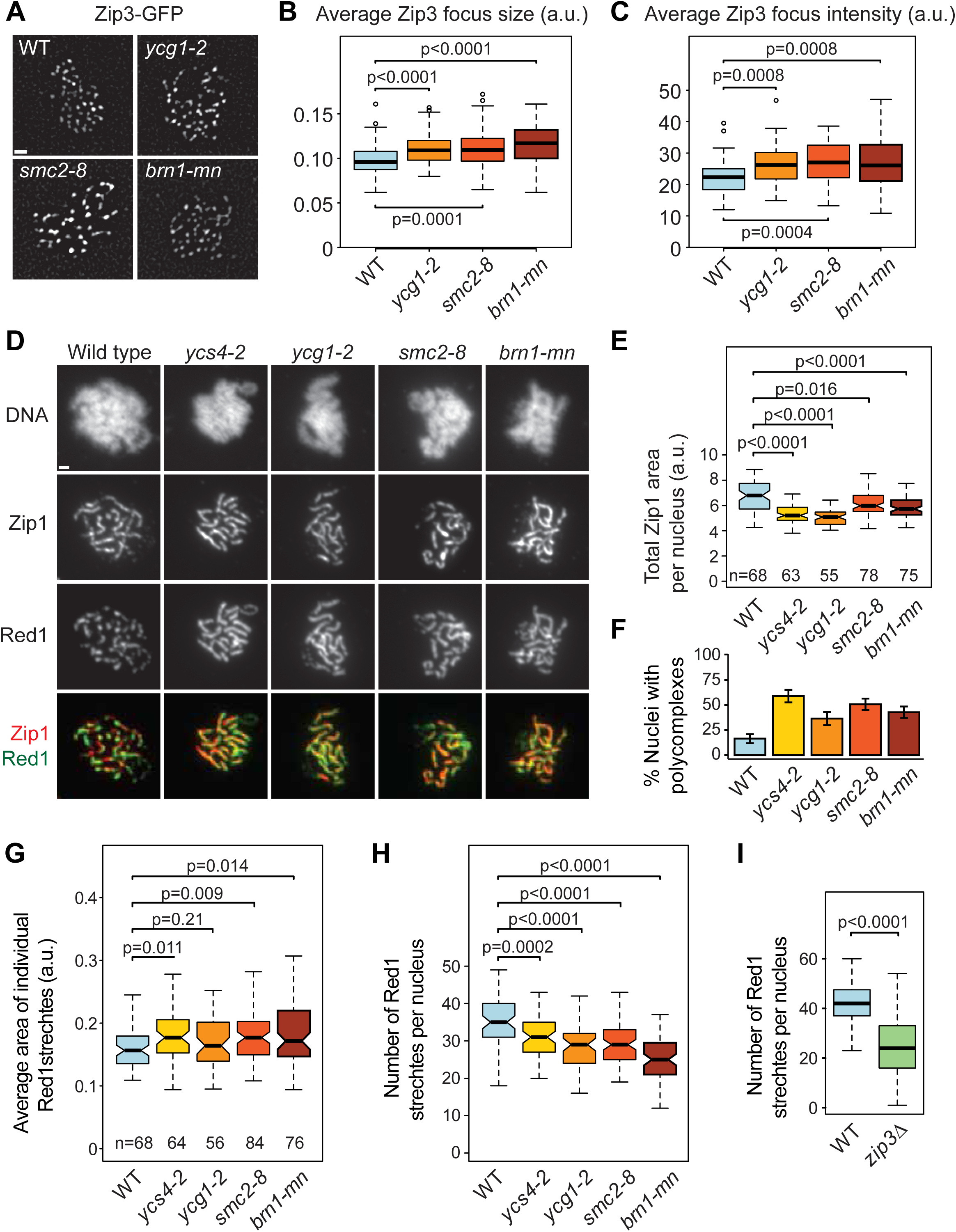
Zip3 foci are larger and meiotic chromosome axes are more continuous in condensin mutants. (**A-C**) Analysis of Zip3-GFP foci at pachynema in wild type [H13116], and *ycg1-2* [H13117], *smc2-8* [H13118], and *brn1-mn* [H13141] mutants at 34℃. (**A**) Representative images of Zip3-GFP in the indicated strains. (**B**) Quantification of the average size of individual Zip3-GFP foci per nucleus in the indicated strains. The number of analyzed nuclei is indicated. (**C**) Quantification of the average signal intensity of individual Zip3-GFP foci in the same nuclei as in (B). Wilcoxon rank sum test with continuity correction. (**D-H**) Analysis of chromosome synapsis (Zip1) and chromosome axes (Red1) at pachynema in wild-type [H7797], *ycs4-2* [H8601], *ycg1-2* [H11281], *smc2-8* [H11364], and *brn1-mn* [H12022] mutants at 34℃. (**D**) Representative images showing the binding of Zip1 (red) and Red1 (green) in the indicated strains. DNA is stained with DAPI. Bar = 1 µm. (**E**) Quantification of total synapsis (Zip1 area) in the indicated strains. The number of analyzed nuclei is indicated. Wilcoxon rank sum test with continuity correction. (**F**) Incidence of Zip1 aggregates (polycomplexes) in the same nuclei as in (E). (**G**) Quantification of the average signal of individual Red1 stretches as a measure of axis fragment length. The number of analyzed nuclei is indicated. (**H**) Quantification of the total number of Red1 stretches per nucleus for the same nuclei as in (G). (**I**) Quantification of the total number of Red1 stretches at pachynema in wild-type [H11471] and *zip3Δ* mutants [H12925]. G-I: Wilcoxon rank sum test with continuity correction. See also **Figure S5**.

Previous work had indicated that condensin mutants are defective in SC formation ^23^. However, those analyses were performed in cells after a prolonged prophase arrest, and the most severe mutant phenotype (*ycs4S*) was later attributed to a genetically linked variant in the promoter of *RED1* ^66^. Therefore, we revisited SC formation in condensin mutants that were allowed to progress through meiosis with normal kinetics. We found that substantial SC formation occurred in all mutants as judged by the incorporation of the transverse filament protein Zip1 (**Figure 5D**), although *ycs4-2* and *ycg1-2* exhibited mildly reduced SC length (**Figure 5E**). In addition, all condensin mutants frequently exhibited Zip1 aggregates (polycomplexes), which are a sensitive indicator of synapsis defects (**Figure 5F**) ^67^. Together, these findings indicate a mild defect in SC formation in the absence of condensin function.

The lateral elements of the SC, composed of Hop1 and Red1, typically exhibit a fragmented or discontinuous staining pattern along synapsed chromosomes ^7,68,69^. Analysis of Red1 staining showed that this fragmentation was significantly reduced in condensin mutants, resulting in longer and fewer Red1 stretches (**Figure 5D, G-H**). These data indicate that condensin is required to create discontinuities along the axes of synapsed chromosomes. In line with this interpretation, *zip3*Δ mutants, which fail to recruit condensin to crossover designated sites, also showed more continuous Red1 stretches (**Figure 5I**). ChIP-seq analysis of Hop1, a binding partner of Red1, showed only minor differences in the levels and chromosomal distribution of Hop1 in condensin mutants (**Figure S6A-B**), consistent with the notion that condensin-dependent axis remodeling occurs locally near crossover sites, which vary among cells.

The increased axial continuity in condensin mutants was associated with a pronounced increase in nuclei containing chromosomal DAPI clouds in a characteristic “parallel track” configuration (**Figure 6A-B**). In wild-type nuclei, this arrangement is usually transient. However, nucleus-wide parallel DAPI tracks are a feature of several chromosome remodeling mutants. These include mutants lacking Pch2, which is responsible for removing Hop1 and Red1 from chromosomes ^7,68^, as well as specific mutants in Zip1 and topoisomerase II (Top2) that are defective or delayed in Pch2 recruitment ^7,70^. Therefore, we asked whether condensin mutants also fail to recruit Pch2. Contrary to this expectation, we observed a significant increase of Pch2 focus intensity on the chromosomes of condensin mutants (**Figure 6C-E**). These findings indicate that the parallel track phenotype can be genetically separated from Pch2 recruitment. Although the increased Pch2 recruitment may hint that Pch2 cannot function properly in the absence of condensin, these data imply that the increased continuity of axial element signals in condensin mutants is not simply due to a failure to recruit Pch2.

**Figure 6.**
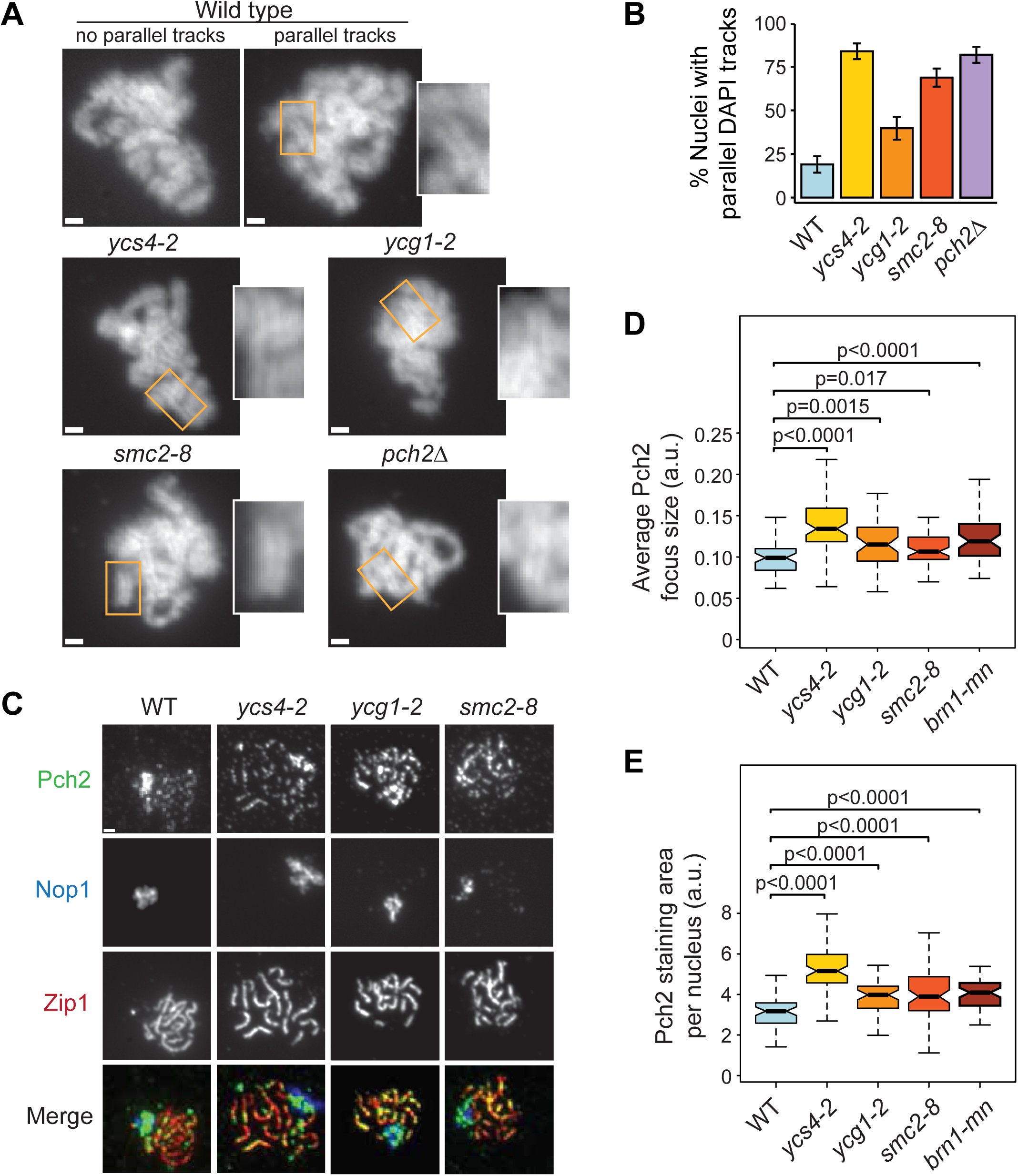
Abnormal chromatin morphology and Pch2 accumulation in condensin mutants. (**A**) Representative images of chromatin morphology of spread pachytene chromosomes strained with DAPI in wild type [H7797], *ycs4-2* [H8601], *ycg1-2* [H11281], and *smc2-8* [H11364] mutants at 34℃. Wild-type examples of nuclei with or without parallel DAPI tracks and *pch2Δ* mutants [H11758], which have strong parallel-track phenotype ^7^, are included for comparison (at 34℃). Bar = 1 µm. Insets highlight examples of parallel DAPI tracks. (**B**) Quantification of nuclei with predominantly parallel DAPI tracks in the indicated strains. (**C**) Representative images showing the relative binding of Pch2 (green) in the nucleolus (marked by staining with Nop1 in blue) and along the SC (marked by Zip1 in green) of spread pachytene chromosomes in wild-type [H7797], *ycs4-2* [H8601], *ycg1-2* [H11281], and *smc2-8* [H11364] and *brn1-mn* [H12022] mutants at 34℃. (**D**) Quantification of average Pch2 focus intensity in the indicated strains. (**E**) Quantification of the average Pch2 staining area per nucleus in the strains analyzed in (D). Wilcoxon rank sum test with continuity correction. See **Figure S6**.

### Condensin reduces cohesin near sites of crossover designation

Condensin was previously shown to promote the chromosomal removal of Rec8-cohesin ^32^, which is the major recruiter of chromosomal Red1 ^43,71,72^. Therefore, the breaks in Red1 continuity could be secondary to a local loss of Rec8 from synapsed chromosomes. In line with these observations, we found that Rec8 signal was consistently more continuous in condensin mutants, resulting in fewer Rec8 stretches (**Figure 7A-B**). Moreover, Rec8 staining at crossover-designated sites, marked by Zip3, was significantly lower in wild-type cells relative to condensin mutants (**Figure S7A-B**). Removal of cohesin from prophase chromosomes requires phosphorylation of Rec8 on serine 521 ^73^. Co-staining of chromosome spreads revealed that foci of phosphorylated Rec8-S521 frequently occurred in close proximity to Zip3 foci (**Figure 7C**). In condensin mutants, the number and intensity of phospho-Rec8-S521 foci was significantly reduced (**Figure 7D-F**), suggesting that condensin regulates Rec8 removal by promoting the formation of phospho-S521-positive chromosomal domains. Together, these data indicate that condensin recruitment to crossover sites is important for restructuring the Rec8-dependent axial element.

**Figure 7.**
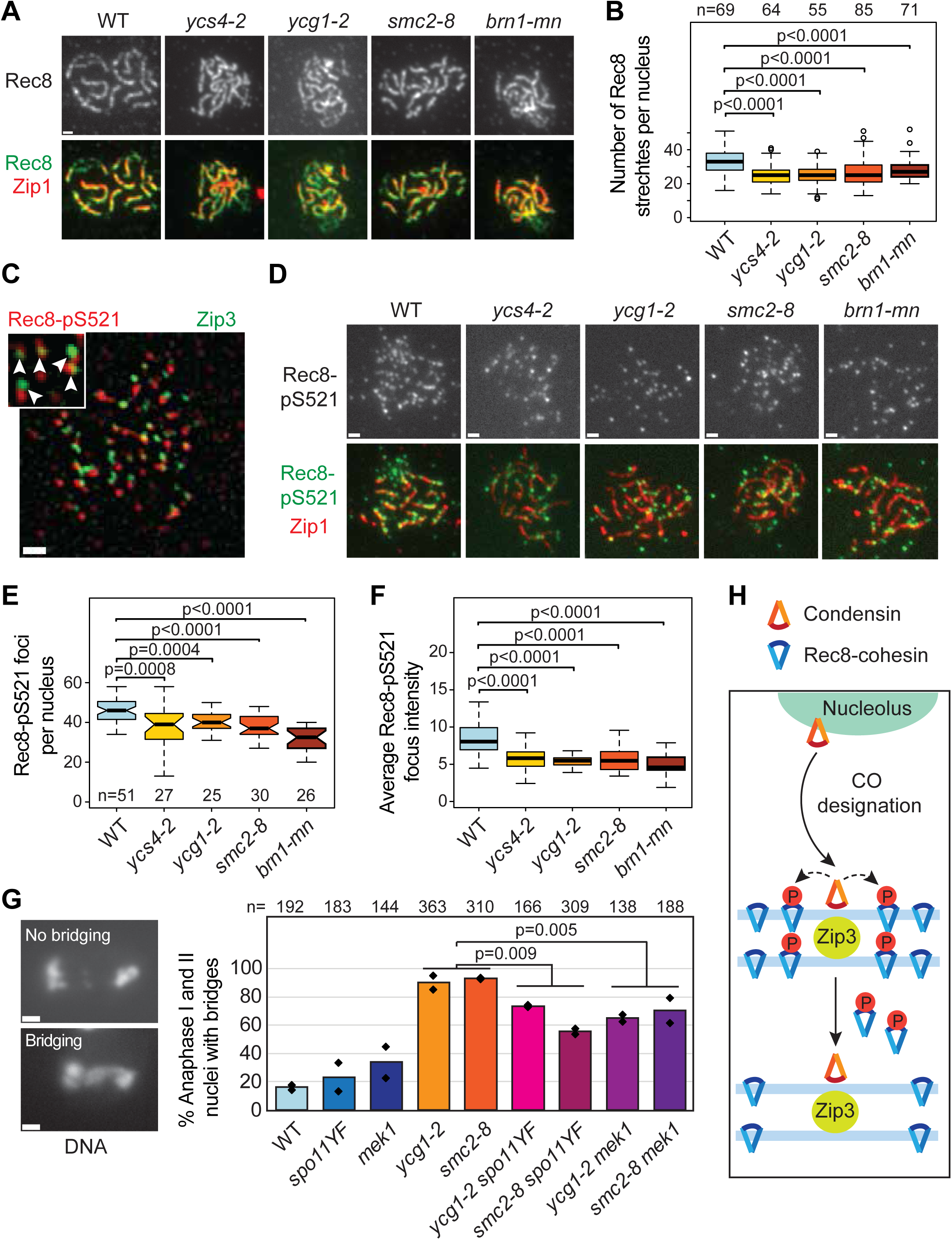
Continuous cohesin axes and interhomolog recombination-dependent meiotic missegregation in condensin mutants. (**A-B**) Analysis of Rec8 binding on spread pachytene chromosomes in wild-type [H7797], *ycs4-2* [H8601], *ycg1-2* [H11281], *smc2-8* [H11364], and *brn1-mn* [H12022] mutants at 34℃. (**A**) Representative images showing the binding of Zip1 (red) and Red1 (green) in the indicated strains. DNA is stained with DAPI. Bar = 1 µm. (**B**) Quantification of the total number of Rec8 stretches per nucleus. The number of analyzed nuclei is indicated. (**C**) Representative image of a wild-type spread nucleus [H13116] showing adjacent foci of Zip3-GFP and phospho-Rec8-S521 (Rec8-pS521). Arrowheads in zoomed-in detail indicate examples of close apposition of foci. (**D**) Representative images of spreads stained for DNA (DAPI), Zip1 (red) and Rec8-pS521 (green) in wild type [H7797], *ycs4-2* [H8601], *ycg1-2* [H11281], *smc2-8* [H11364], and *brn1-mn* [H12022] mutants at 34℃. (**E**) Quantification of the total number of Rec8-pS521 foci per nucleus in the indicated strains. The number of analyzed nuclei is indicated. (**F**) Average focus intensity of the data shown in (D). Wilcoxon rank sum test with continuity correction. (**G**) Quantification of chromosome bridging in anaphase I and II nuclei in wild type [H7797], *ycg1-2* [H11281], *smc2-8* [H11364], *spo11-YF* [H12173], *ycg1-2 spo11-YF* [H12227], *smc2-8 spo11-YF* [H12245], *mek1Δ* [H12137], *ycg1-2 mek1Δ* [H12194], *smc2-8 mek1Δ* [H12205]. The panels on the left show examples of spread nuclei with normal meiosis I chromosome separation or anaphase I chromosome bridging. DNA is stained with DAPI. The number of analyzed nuclei is indicated. Welch two-sample *t*-tests. (**H**) Schematic summarizing the proposed regulation of the meiotic chromosome axis by condensin. Upon crossover designation, condensin relocalizes from the nucleolus to the vicinity of crossover repairs sites. There, it promotes the phosphorylation of Rec8 on S521 (red circles) and local removal of cohesin. See also **Figure S7**.

### Crossover-dependent chromosome bridging in condensin mutants

Chromosome bridging is a well-established phenotype of condensin mutants during mitotic cell division ^20^. Intriguingly, bridging during meiotic anaphase I and II depends partly on meiotic DSB formation ^23,32^ (**Figure 7G**), indicating that some of the bridging is a consequence of meiotic recombination. One potential explanation is that chromosome bridging arises from defects in resolving double Holliday junction intermediates into crossovers. However, double Holliday junction resolution occurred normally in condensin mutants (**Figure 4C, G**), indicating that the bridges are not caused by abnormal joint molecules.

Nevertheless, the localization of condensin at sites of crossover repair suggested that the chromosome bridging may be linked to crossover formation. To test this possibility, we deleted the meiosis-specific checkpoint kinase Mek1. In *mek1*Δ, DSBs form at wild-type levels but are primarily repaired using the sister chromatid as a template, leading to a profound drop in recombination between homologous chromosomes ^74,75^. We found that the deletion of *MEK1* reduced chromosome bridging in condensin mutants similarly to *spo11* mutants (**Figure 7G**), indicating that part of the bridging phenotype arises from interhomolog recombination. Therefore, we propose that the localized condensin-mediated removal of cohesin around sites of crossover formation is essential for the separation of homologous chromosomes during the first meiotic division.

## Discussion

Here, we demonstrate that condensin is recruited to designated crossover repair sites to regulate the structure of the meiotic chromosome axis. We propose that the function of meiotic condensin is to promote crossover-associated axis remodeling, to ensure the orderly separation of recombinant chromosomes during meiosis I.

Our data paint a dynamic picture of condensin localization that is in line with available data. Although condensin functions as a loop extruder, condensin tends to be more important for the timely remodeling of chromosome architecture than the stable formation of chromatin loops. Many of condensin’s roles align with this interpretation. For instance, during mitotic anaphase, condensin acutely relocalizes from centromeres to chromosome arms to facilitate the resolution of sister chromatid entanglements ^76,77^, and in meiosis, it transitions from the nucleolus to meiotic centromeres to promote the co-orientation of sister kinetochores ^65^. In addition, yeast condensin compacts and segregates rDNA, clusters tRNA loci, and organizes a topologically associated domain specifically during anaphase ^19,20^. Our findings suggest that some of these activities also take place concurrently with meiotic recombination, perhaps explaining why a certain number of condensin foci are always detectable on meiotic chromosomes. However, overlaid over these background activities, we observed extensive redistribution of condensin from the nucleolus to sites of crossover formation.

The mechanism that directs the recruitment of condensin to crossover sites remains to be determined. One possibility is that components of the crossover machinery, such as the ZMM factors, directly bind condensin, thus coupling the assembly of crossover repair sites to the recruitment of condensin. This mechanism would be akin to the distinct chromosomal receptor motifs that mediate condensin recruitment to pericentromeres and the rDNA ^78^. However, as condensin foci do not directly overlap with Zip3 and Msh4 foci, condensin may also recognize specific chromatin features that are unique to crossover repair intermediates. Condensin preferentially binds single-stranded DNA *in vitro* and can remove the single-stranded binding protein RPA ^79,80^, raising the possibility that RPA-coated D-loops - a feature of crossover intermediates ^81^ - could directly recruit condensin to crossover sites. Alternatively, condensin may be recruited by crossover-associated chromatin, analogous to its recruitment to chromosome arms by histones H2A and H2A.Z during mitotic anaphase ^77^. Although chromatin features unique to crossover sites have not yet been identified, several histone modifications are specifically enriched during meiotic prophase ^82–84^ and could conceivably serve as recruitment signals. Another possibility is that condensin may also be directly recruited by Rec8-cohesin. In line with this possibility, meiosis-specific binding of condensin to chromosomes is largely abolished in *rec8* mutants. A tight interaction of condensin with a larger crossover-associated intermediate or chromatin domain may also explain why successful ChIP-seq analysis of condensin required deproteination prior to re-sonication.

Despite the close association of condensin with crossover repair factors, we find no evidence that condensin directly regulates crossover repair. Crossover levels remain largely unchanged and no aberrant multi-chromatid structures were detected at the well-characterized *HIS4LEU2* and *ERG1* hotspots. Moreover, crossover spacing (interference) was previously reported to be unaffected by condensin mutations ^61^. These findings indicate that condensin is neither a pro-crossover ZMM factor ^54^ nor an anti-crossover factor, like the Sgs1 helicase ^55,85^. Instead, the most prominent defects we observed were related to the aberrant structure of the meiotic chromosome axis, suggesting that condensin is primarily important for the axis reorganization near crossover sites.

Axis reorganization near crossover sites may be important for proper chromosome segregation for a couple of reasons. Crossover recombination can give rise to chromosomal interlocks, where chromosomes become trapped between pairs of synapsing chromosomes. If unresolved, these interlocks can, in rare cases, lead to aberrant chromosome alignment on the metaphase plate and result in mis-segregation ^86^. Interlocks are abundant in mid-prophase but are largely resolved by late prophase ^87,88^, perhaps because of condensin activity. The timing of condensin recruitment to chromosomes is consistent with this hypothesis, as is the observation that interlock removal in *Arabidopsis thaliana* requires topoisomerase II ^89^, which often collaborates with condensin to promote chromosome organization. However, in species like budding yeast, with relatively short chromosomes, interlocks are rare ^17,90^ and cannot fully explain the scale of crossover-dependent chromosome bridging observed in condensin mutants.

Based on these considerations, we favor a model in which condensin is necessary to remove crossover-associated cohesin domains (**Figure 7H**). In organisms with suitable cytology, including grasshoppers and plants, local dissolution of sister chromatid cohesion around crossover sites is readily observable ^14–16^, and in *Sordaria* and rat, Rec8 is locally depleted at crossovers sites ^17,18^. Overall condensin-dependent depletion of Rec8 from late meiotic prophase chromosomes has been reported in budding yeast ^32,73^, and our observations support the model that this removal is linked to crossover repair. Interestingly, the pool of cohesin regulated by condensin is less sensitive to cleavage by separase ^32^. Thus, the processes linked to crossover formation may create separase-resistant cohesin domains, requiring a specialized, separase-independent mechanism of removal.

How condensin mediates cohesin removal remains to be determined. In principle, removal may occur through direct competition, as the loop forming activities of condensin and cohesin influence each other when present on the same chromatin substrate ^91^. Although our experiments did not directly assess the loop-extruding activity of condensin, we note that the *smc2-8* mutation is not a protein null mutation and permits abundant binding of condensin to meiotic chromosomes (**Figure S3D**). This temperature-sensitive mutation, which is caused by an internal epitope tag ^92^, likely interferes with a more specific aspect of condensin function. Supporting this notion, the removal of cohesin during meiotic prophase also requires phosphorylation of Rec8 at Serine 521 by Dbf4-dependent kinase (DDK) ^73,93^. We show that Rec8-pS521 is spatially enriched at crossover sites and that both number and intensity levels of Rec8-pS521 are significantly reduced in condensin mutants. These findings suggest that condensin recruitment to crossover sites helps establish a chromatin environment that promotes the phosphorylation of Rec8, ultimately helping with its timely removal from chiasmata. Given that chromosome bridging during meiosis I is a widely conserved phenotype of condensin mutants across diverse organisms ^33,34,94,95^, similar crossover-associated barriers to chromosome segregation may also need to be resolved in other organisms, including humans.

## Supporting information

Key resources table

## Resource Availability

### Lead contact

Requests for further information and resources should be directed to and will be fulfilled by the lead contact, Andreas Hochwagen.

## Materials Availability

Yeast strains generated in this study are available from the lead contact without restrictions.

## Data and Code Availability

--**Data**

Meiotic ChIP-seq data have been deposited at the Gene Expression Omnibus (GEO) as GSE154722 (‘https://www.ncbi.nlm.nih.gov/geo/query/acc.cgi?acc=GSE154722’) and are publicly available as of the date of publication. The analysis of condensin in asynchronous cells used previously published ChIP-seq data: GEO accession GSE106104. For ranking hotspots based on Spo11-oligos, we the used publicly available GEO accession GSE48299. Red1 profiles were taken from the publicly available GEO accession GSE70112.

--**Code**

All original code has been deposited at Zenodo and is publicly available. Computer scripts used for processing Illumina reads are available at: https://doi.org/10.5281/zenodo.16783225

Computer scripts used for data analysis are available at: https://doi.org/10.5281/zenodo.16783262

**--Additional Information**

Any additional information required to reanalyze the data reported in this paper is available from the lead contact upon request.

## Acknowledgements

We thank F. Uhlmann, P. Jordan, N. Kleckner, H. Yu, and A. Strunnikov for sharing strains, N. Hollingsworth, A. Shinohara, and G. Brar for sharing antibodies, and H. Murakami for helpful discussions. This work was supported in part through the NYU IT High Performance Computing resources, services, and staff expertise. We acknowledge the Zegar Family Foundation for their support. We thank the NYU Center for Genomics and Systems Biology Genomics Core for their assistance and resources. This work was supported by National Institutes of Health grant R35 GM148223 to AH and National Research Foundation of Korea (NRF) grant funded by the Korean government (MSIT) [RS-2023-00208191 to KPK]. ARR acknowledges support from a Fleur Strand Graduate Fellowship from the Department of Biology, as well as a Henry MacCracken Fellowship and a Dean’s Dissertation Fellowship from the NYU Graduate School of Arts and Science. The funders had no role in the preparation of this manuscript.

## Author contributions

Conceptualization, V.A.L, T.E.M, and A.H.; Investigation, V.A.L, T.E.M., S.H., A.R.R., J.H., K.P.K. and A.H.; Software, T.E.M.; Formal Analysis, V.A.L, T.E.M., S.H., A.R.R., J.H., K.P.K.; Writing – Original Draft, V.A.L., T.E.M, and A.H.; Writing – Review & Editing, V.A.L, T.E.M., S.H., A.R.R., J.H., K.P.K. and A.H.

## Declaration of Interest

The authors declare no conflict of interest.

## STAR Methods

### EXPERIMENTAL MODEL AND STUDY PARTICIPANT DETAILS

All strains used in this study are isogenic or congenic with the *S. cerevisiae* SK1 background. See also **Table S1**. Gene disruption and epitope tagging were carried out using a PCR-based protocol ^96^. The Zip3-13myc and Zip3-GFP tags were provided by N. Kleckner and P. Jordan, respectively ^61,97^. The Smc4-PK9 tag was transferred from W303 ^36^ using allele replacement by transformation. The Smc4-GFP tag is from ^37^ and the *pCLB2-BRN1* (*brn1-mn*) allele was described in ^65^. The temperature-sensitive condensin alleles were introduced into SK1 by backcrossing a minimum of 7 times. The *ycg1-2* and *ycs4-2* alleles are from ^23^, *smc2-8* and *smc4-1* are from ^98^. *smc2-8* is the result of an epitope tag within Smc2, as previously noted by ^92^. To induce synchronous meiosis, strains were inoculated in BYTA for 16.5 hours at 30°C, washed two times with water and resuspended at an OD_600_ = 2.0 at 30°C in SPO medium ^99^. Temperature sensitive strains as well as the wild-type control and the *brn1-mn* strain, were inoculated at OD_600_ = 0.3 in BYTA medium for 18 hours at 25°C and were inoculated in SPO medium at an OD_600_ = 2.0 at 25 °C for 1 hour before shifting to 34°C. For the experiments in **Figure 4** and **Figure S4**, cells were pre-grown in YPA and transferred into SPM at 25°C and shifted to 34°C at 2.5h ^100,101^. The later temperature shift in these experiments is based on the slower meiotic kinetics under YPA/SPM culture conditions. For all time courses, efficient meiotic entry was confirmed by monitoring DNA content using flow cytometry. Only cultures that efficiently entered meiosis were analyzed. Flow cytometry was also used to orient the timing of temperature shifts under different culture conditions.

## METHOD DETAILS

### ChIP-seq analysis

Chromatin immunoprecipitation (ChIP) was performed as described ^99^. For all ChIP-seq samples, 25 mL meiotic culture were collected 3h after meiotic induction. Samples were immunoprecipitated with 2 μL anti-Red1 serum (Lot#16440, kind gift of N. Hollingsworth ^102^), 2 μL anti-Hop1 serum (kind gift N. Hollingsworth ^103^), 20 μL anti-V5 (PK9) agarose affinity gel (Sigma), or 10μL of anti-HA 3F10 antibody (Roche Applied Science) per IP. Library preparation was performed using Illumina TruSeq DNA Sample Prep Kits v1, but adapters were used at 1:20 dilution. Library quality was confirmed by Qubit HS assay kit and Agilent 2200 TapeStation. 150-bp sequencing was accomplished on an Illumina HiSeq 2500 or NextSeq 500 instrument. ChIP-seq data are averages of at least two biological replicates collected from samples grown on separate days and sequenced independently. The only experimental conditions with a single replicate were the no-tag control under traditional ChIP conditions and the ChIP of Hop1.

### PK9-specific adjustments to ChIP-seq protocol

Many known condensin binding sites map to nucleosome-free regions or otherwise open chromatin ^36,92^. These regions are more prone to breakage by sonication and can create background signals independent of enrichment by ChIP ^104^. In addition, our benchmarking experiments indicated that PK9 antibody preferentially binds to these regions independently of the PK9 tag. To account for these non-specific signals, we included no-tag controls with all ChIP-seq analyses of Smc4-PK9. In addition, to increase the concentration of reads from longer ChIPed fragments, ChIP and input DNA were re-sonicated using a Bioruptor Pico (Diagenode, NJ, USA) with the following settings: 30 seconds ON and 30 seconds OFF for 5 cycles. This change to the general ChIP-seq protocol is not novel ^105^.

### Chromosome spreads

Meiotic nuclear spreads were performed as described ^106^. GFP was detected using a polyclonal chicken anti-GFP antibody (abcam13970) at 1:100 dilution and an Alexa Fluor 488-conjugated anti-chicken secondary antibody at 1:200 dilution. Zip1 was detected using a polyclonal goat antibody against a C-terminal peptide of Zip1 (yC-19; Santa Cruz Biotechnology) at 1:200 and a Cy3-conjugated anti-goat secondary antibody at 1:200. Red1 was detected using rabbit serum (Lot#16440, kind gift of N. Hollingsworth ^102^) at 1:100 and Alexa 488-conjugated anti-rabbit secondary antibody at 1:200. Nop1 was detected using a mouse monoclonal Nop1 antibody (EnCor Biotechnology) at 1:400 and a Cy5-conjugated anti-mouse secondary antibody at 1:200. Rec8 was detected with a guinea pig anti-Rec8 serum (kind gift of N. Hollingsworth ^107^) at 1:400 and a Cy5-conjugated anti-guinea pig secondary antibody at 1:200. Pch2 was detected by using rabbit anti-Pch2 serum (kind gift from A. Shinohara ^7^) and Alexa 488-conjugated anti-rabbit secondary antibody at 1:200. Myc-tagged proteins were detected using a mouse monoclonal anti-myc antibody (4A6; Millipore Sigma) and a Cy5-conjugated anti-mouse secondary antibody at 1:200. Phospho-Rec8-521 was detected with a phospho-specific rabbit antiserum (kind gift of G. Brar ^108^) at 1:100 and Cy5- or Alexa 488-conjugated anti-rabbit secondary antibodies at 1:200. Fpr3 was detected by using rabbit anti-Fpr3 serum (kind gift from J. Thorner ^109^) at 1:400 and Cy3-conjugated anti-rabbit secondary antibody at 1:200. All secondary antibodies came from Jackson ImmunoResearch. Images were obtained on a Deltavision Elite imaging system (GE Applied Precision) adapted to an Olympus IX 17 microscope.

### Whole-cell immunofluorescence of spindles

A total of 300 μL meiotic culture was collected, pelleted, and fixed in formaldehyde as in ^66^. Permeabilized cells were stained with monoclonal rat anti-tubulin antibody (YOL1/34; Santa Cruz Biotechnology) at 1:100 dilution and an Alexa Fluor 488-conjugated anti-rat secondary antibody (Jackson ImmunoResearch) at 1:200. DNA was visualized using DAPI mounting medium (VectaShield).

### Protein extraction and western blotting

At the indicated time points, 5 mL of meiotic culture were harvested and resuspended in 5% trichloroacetic acid (TCA), followed by incubation on ice for at least 10 minutes. Samples were then washed with 1X TE buffer and resuspended in 1 M unbuffered Tris, pH ∼11, for 2 minutes at room temperature. Cells were lysed in lysis buffer (10 mM Tris-HCl, pH 7.5, 1 mM EDTA, 2.75 mM dithiothreitol). After lysis, 5X SDS sample buffer was added, and samples were incubated at 100°C for 2 minutes before analysis. Protein samples were separated on an 8–12% stain-free SDS-PAGE gel (Bio-Rad) and transferred onto a PVDF membrane. Membranes were stained with Ponceau S to verify transfer efficiency. The Ycs4–13Myc fusion protein was detected using a mouse monoclonal anti-Myc antibody (clone 4A6, Millipore) at a dilution of 1:1,000 and an HRP-conjugated anti-mouse secondary antibody (Kindle Biosciences) at 1:1,000.

### Pulsed-field gel electrophoresis and Southern analysis of chromosome III

For every time point, 10 mL of meiotic culture were collected and killed by adding sodium azide (0.1% final). Agarose plugs were prepared as in ^110^. Plugs were melted and loaded into a 1.1% Seakem LE agarose gel and run in 0.5X TBE on CHEF Drive II apparatus (BioRad) using a 5 to 30 second ramp, run time 38h, voltage 5.4V/cm. After the run the DNA was stained with ethidium bromide and exposed to UV light for 4 minutes to fragment DNA. DNA was depurinated in 0.25 M HCl for 30 mins and denatured in 0.4M NaOH for 40 mins before transferring onto a HybondXL membrane (Cytiva) in 0.4M NaOH/0.6M NaCl overnight. The membrane was probed with and alpha-^32^P-dCTP-labeled probe using a probe sequence near *MRC1* on chromosome III ^110^. The membrane was exposed on Fujifilm imaging plates and imaged using a Typhoon FLA9000 instrument.

### Analysis of recombination intermediates

Genomic DNA preparation and physical analysis were performed as described previously ^64,100,101^. To synchronize yeast cell cultures, overnight YPD-cultured cells were transferred to SPS medium (1% potassium acetate, 1% bacteriological peptone, 0.5% yeast extract, 0.17% yeast nitrogen base with ammonium sulfate and without amino acids, 0.5% ammonium sulfate, 0.05 M potassium biphthalate, adjusted to pH 5.5 with 10 N KOH) and grown for 18 h at 25℃. Meiosis was induced by transferring the SPS-cultured cells to pre-warmed SPM medium (1% potassium acetate, 0.02% raffinose, two drops of Antifoam 204). SPM-cultured cells were harvested at 0, 2.5, 3.5, 4, 5, 6, 7, 8, 10 and 24 h after being transferred to the SPM medium and then cross-linked with psoralen under ultraviolet light at a wavelength of 360 nm. For the analysis of meiotic cells grown at 34℃, cells were cultured at 25°C for up to 2.5 h and then shifted to 34°C in SPM medium. For 1D and interhomolog crossover (CO)/interhomolog noncrossover (NCO) gel analysis, 2 μg of genomic DNA was digested with XhoI and NgoMIV restriction enzymes. The digested DNA samples were loaded onto a 0.6% UltraKem LE agarose gel (Young Science, Y50004) in 1X TBE buffer and subjected to electrophoresis at ∼2 V/cm for 24 hours. The DNA in the gel was stained with 0.5 μg/ml ethidium bromide (EtBr) for 30 minutes. For 2D gel analysis, 2.5 μg of genomic DNA was digested with XhoI and loaded onto a 0.4% Seakem Gold agarose gel (Lonza) in 1X TBE buffer. Electrophoresis was performed at ∼1 V/cm for 21 hours. The gel was then stained with 0.5 μg/ml EtBr. For the second-dimension electrophoresis, gel strips were excised and placed onto a 2D gel tray, overlaid with 0.8% UltraKem LE agarose gel containing EtBr. Electrophoresis was conducted at ∼6 V/cm at 4°C for 6 hours. For Southern blot analysis, hybridization for *HIS4-LEU2* was performed using ‘Probe A’, which was labeled with ³²P-dCTP using a random primer DNA labeling kit (Enzynomics). Southern blot analysis of the *ERG1* locus was performed as described in ^64^.

## QUANTIFICATION AND STATISTICAL ANALYSIS

### ChIP-seq data analysis

Sequencing reads were mapped to the SK1 genome ^111^ using Bowtie ^112^ and Bowtie2 ^113^. Reads that mapped to only one location without mismatches were used in further analyses. Further processing was completed using MACS-2.1.0 (https://github.com/taoliu/MACS) ^114^. Single-end reads were extended towards 3’ ends to a final length of 200 bp and probabilistically determined PCR duplicates were removed. Pileups of both the input and ChIP libraries were SPMR-normalized (signal per million reads), followed by a calculation of the fold-enrichment of the ChIP data over the input data. Before plotting, all data was normalized to produce a genome average of 1 to allow for some comparability between experiments. When available, reads of biological replicates were combined prior to MACS2 analysis. For analysis of the rDNA, reads were mapped to a single rDNA repeat using Bowtie with default settings. The 95% confidence intervals were calculated by bootstrap resampling from the data 1,000 times with replacement.

### Analysis of immunofluorescence signals

Spread images were analyzed using softWoRx 5.0 and custom scripts in Fiji. Statistical significance was tested in R using either a Wilcox rank sum test with continuity correction or a Welch’s two-sample t-test, as indicated in the figure legends. Where appropriate, p values were corrected for multiple hypothesis testing using the Benjamini-Hochberg correction. For analysis of chromosome bridging, DAPI-stained nuclear masses were evaluated for the presence or absence of connections between masses as described in Yu et al. ^23^. Briefly, we counted DAPI-stained nuclei that exhibited individually defined masses and used Tubulin staining to stage whole-cells in anaphase I or anaphase II. Counts from both stages were added together. Masses that were bridged, meaning connected by a DAPI signal, were scored as bridging. Masses that were fully separated and showed no DAPI signal between them were counted as separated in the analysis.

### Colocalization analysis

Colocalization was analyzed in Fiji by utilizing the DiAna plugin for 3D localization and distance analysis ^115^. A mask was created from the DAPI channel and regions of the DAPI signal that did not overlap with Zip1 were subtracted to exclude nucleolar condensin signals. This step was necessary because neither Zip3 nor Msh4 typically localizes within the nucleolus. Next, we processed the images with the DiAna plugin to quantify center-to-center distances between foci. To assess statistical spatial robustness, we performed a shuffle analysis with 100 iterations using the DiAna plugin, generating a uniform distribution representing predicted distances. The median focus distances per nucleus for the experimental and randomized datasets were used for statistical analysis. For the overlay images showing colocalization (Figures 3A-B [bottom right panels] and 7C), foci were sharpened using Fiji’s “Blur” function, followed by image subtraction.

### Southern blot quantification

Hybridization signals corresponding to Mom, Dad, DSBs, COs, SEIs, and dHJs were visualized using a phosphorimager and quantified using the Bio-Rad Quantity One software.

## Notes

### Competing Interest Statement

The authors have declared no competing interest.

### Summary of Updates

Figure 2, Supplemental Figure 2, and Figure 7 revised to include additional data; text has been revised to describe that data and to expand description of statistical analyses

