## Supplemental Figures and Material for "Crossover designation recruits condensin to reorganize the meiotic chromosome axis"

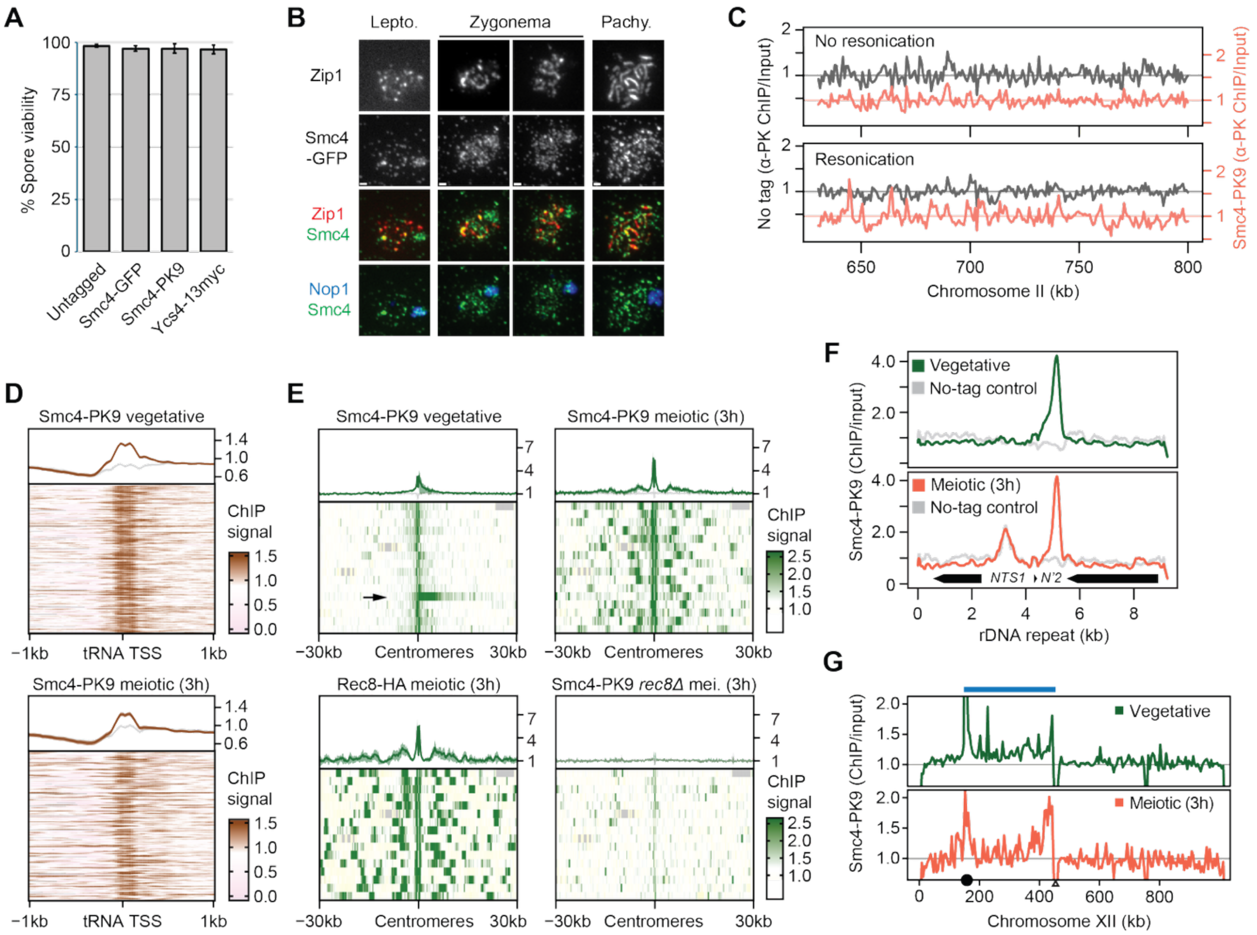


**Figure S1. Condensin tags and chromosomal localization. Related to Figure 1.** (**A**) Spore viability analysis of condensin-tagged strains used in this study. Approximately 100 tetrads were dissected from untagged cells [H7797] and strains carrying homozygous Smc4-GFP [H11471], Smc4-PK9 [H6408], or Ycs4-13myc [H9077] tags. (**B**) Representative images of surface-spread chromosomes from wild-type nuclei [H11471] showing the redistribution of Smc4-GFP (green) from the nucleolus (marked by Nop1, blue) to chromosome arms upon synapsis initiation (marked by Zip1, red). (**C**) ChIP-seq binding profiles of Smc4-PK9 (orange) [H6408] and no-tag control (gray) [H7797] over a representative region of chromosome II before and after re-sonication. (**D**) Enrichment of Smc4-PK9 at tRNA genes in wild-type vegetative and meiotic prophase samples. ChIP-seq values were averaged over 25-bp windows around tRNA transcription start sites (TSS) across a 1-kb window. Rows sorted by tRNA size. Brown is high enrichment; pale pink is depletion. Mean and 95% confidence intervals are shown in line graph directly above heatmap. Gray line indicates mean and 95% confidence intervals of the no-tag control. (**E**) Heatmaps of Smc4-PK9 enrichment around the 16 pericentromeres from vegetative cells [H6408], in meiotic prophase (t=3h) [H6408], and in meiotic *rec8*Δ mutants [H7660]. Rec8-HA enrichment in meiotic prophase (t=3h) [H4471] is shown for comparison. Arrow indicates pronounced condensin domain reaching from *CEN12* toward the rDNA in vegetative cells. (**F**) Smc4-PK9 binding at a single average rDNA repeat in vegetative cells (green) and cells in meiotic prophase (orange). Gray lines are no-tag controls. Black arrows indicate the positions of three rRNA coding regions: *RDN18*, *RDN5*, and *RDN25* from left to right. (**G**) Smc4-PK9 binding pattern on chromosome XII in wild-type vegetative (green) and meiotic prophase cells (orange). Black circle marks the centromere, triangle indicates the rDNA, blue bar denotes the interstitial region between the *CEN12* and the rDNA. Read counts over the rDNA array were set to zero, as no reads were mapped to that region. Gray horizontal bar represents genome-wide average.

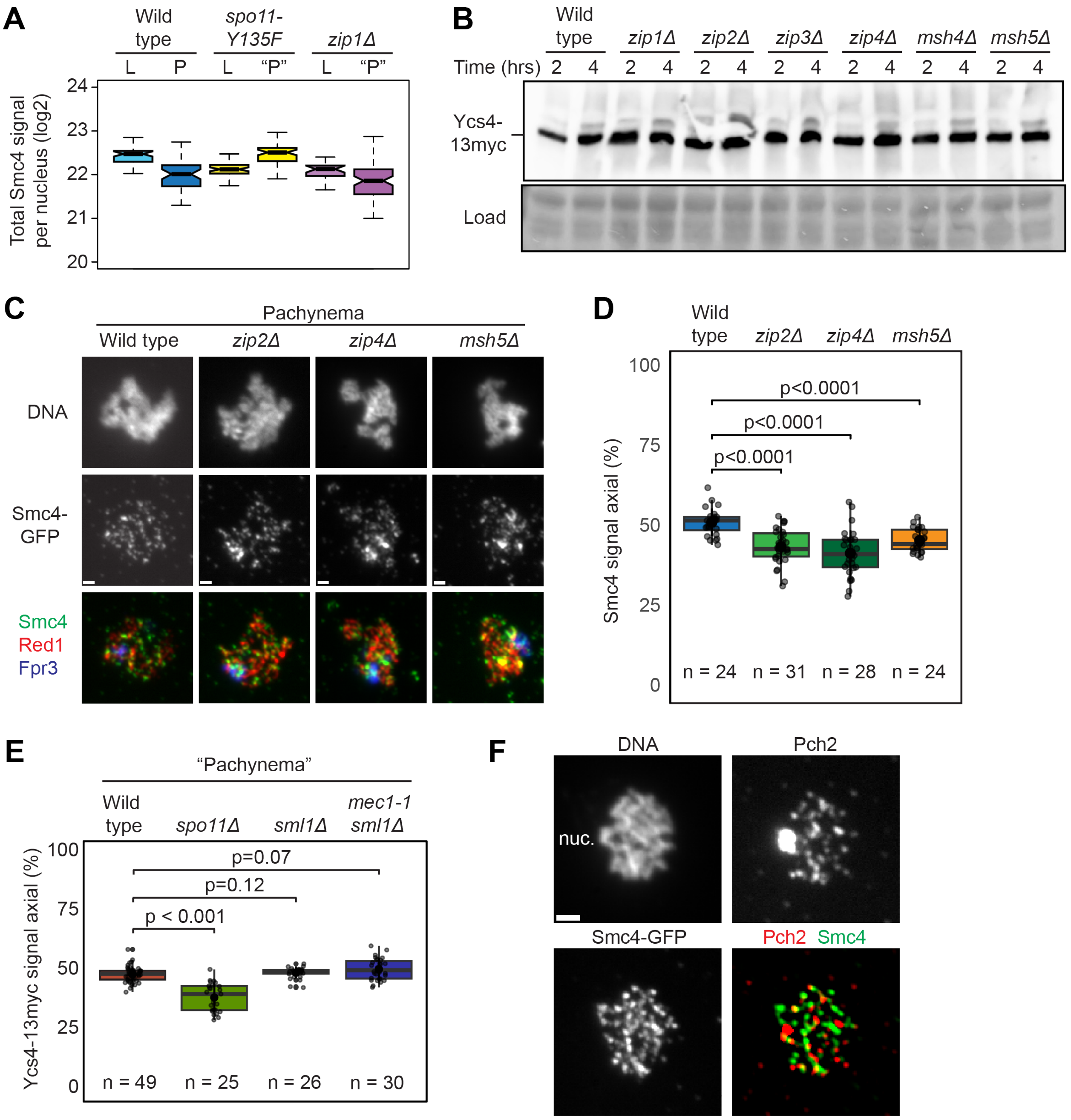


**Figure S2. Effects of recombination regulators on condensin recruitment. Related to Figure 2.**

(**A**) Quantification of total Smc4-GFP fluorescence per surface-spread nuclei in wild-type [H11471], *spo11-YF* [H9441], and *zip1*Δ [H11499] strains as in Figure 2A. Nuclei were staged as leptonema (L) or pachynema (P) based on Red1 staining. In *spo11-YF* and *zip1*Δ mutants, continuous Red1 staining was classified as pachynema-like ("P"). (**B**) Western blot analysis of Ycs4-13myc protein as a proxy for condensin levels in different *zmm* mutants. Despite multiple attempts, we were unable to detect reliable western signals for Smc4-GFP with our available antibodies, possibly due to epitope denaturation by the detergent. Samples were collected from synchronous meiotic cultures of wild type [H13391], *zip1*Δ [H13373], *zip2*Δ [H13376], *zip3*Δ [H13394], *zip4*Δ [H13379], *msh4*Δ [H13382], and *msh5*Δ [H13382] at 2 and 4 hours after meiotic induction, corresponding to early and late prophase, respectively. Ponceau S staining confirmed equal protein loading across samples. (**C**) Representative images showing Smc4-GFP (green) distribution on surface-spread nuclei from wild type [H11471], and *zip2*Δ [H13167], *zip4*Δ [H13307], and *msh5*Δ mutants [H13306]. Chromosomes were co-stained for DNA (DAPI), Red1 (red), and the nucleolar marker Nop1 (blue). Nuclei were staged based on Red1 staining: punctate Red1 staining was classified as leptonema and individualized Red1 axes were classified as pachynema. Bar = 1 µm. (**D**) Smc4-GFP signal in pachynema was quantified based on overlap with Nop1 and classified as nucleolar, while non-overlapping signal with Nop1 was classified as axial. The percentage of axial signal relative to total signal is shown. Wilcoxon rank sum test with continuity correction. (**E**) Quantification of Ycs4-13myc signal in pachynema (or pseudo-pachynema) surface-spread nuclei from wild-type [H9077], *spo11*Δ [H11058], *sml1*Δ [H13208], and *mec1-1 sml1*Δ [H11014] strains. Signal overlapping with the nucleolar marker Fpr3 was classified as nucleolar, and non-overlapping signal was classified as axial. The percentage of axial signal relative to total signal is shown. The number of nuclei analyzed is indicated. Wilcoxon rank sum test with continuity correction. (**F**) Representative images showing the relative distribution of Smc4-GFP (green) compared to Pch2 (red) on a surface-spread pachytene nucleus of a wild-type cell [H11471]. DNA was stained with DAPI. nuc. = nucleolus, as apparent from the unsynapsed chromatin.

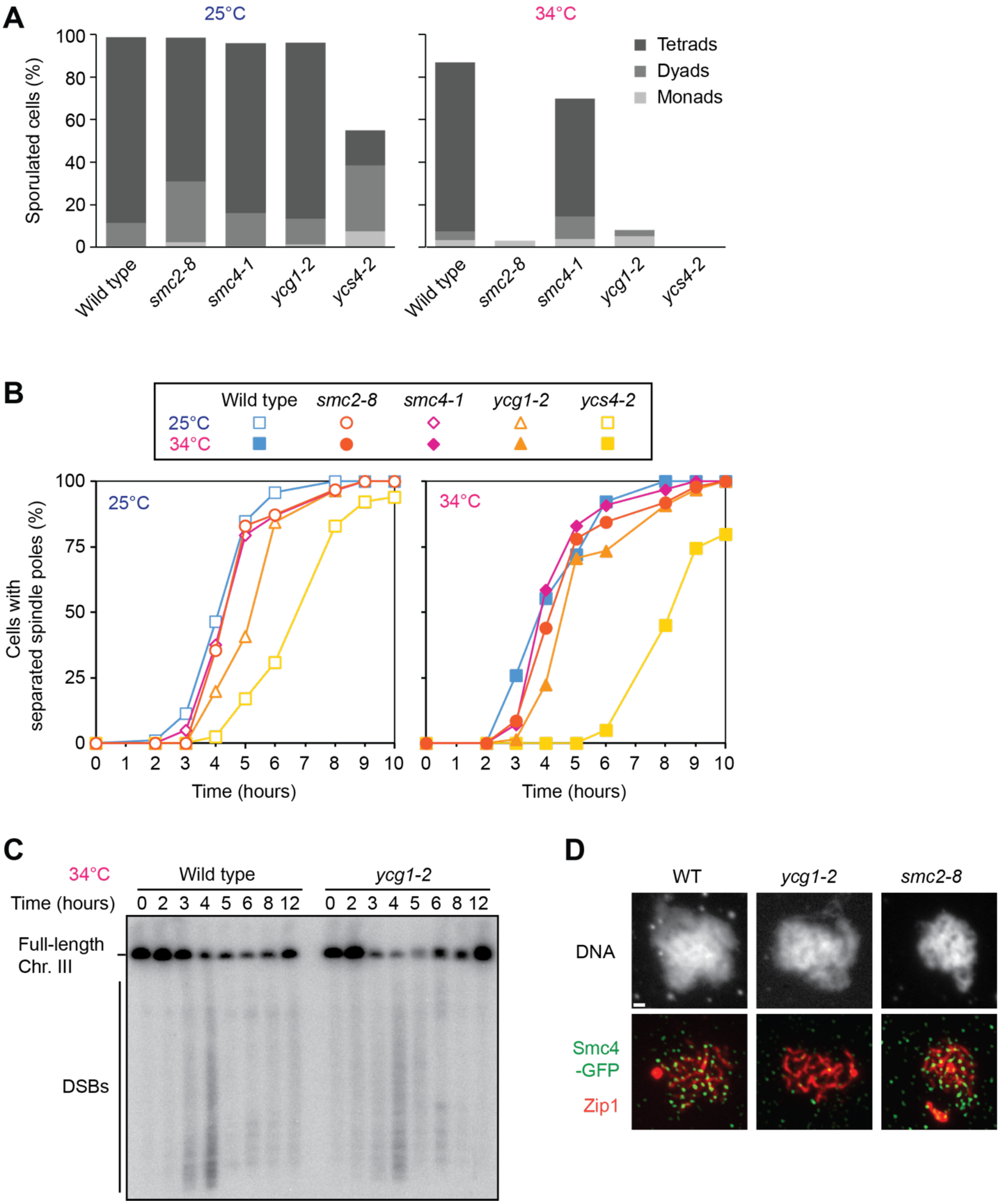


**Figure S3. Condensin temperature-sensitive mutants have sporulation defects but form DSBs and meiotic spindles with largely normal kinetics. Related to Figures 4-7.**
(**A**) Quantification of spore formation of wild-type [H7797], *ycs4-2* [H8601], *ycg1-2* [H11281], *smc4-1* [H11363], and *smc2-8* [H11364] strains incubated at 25°C and 34°C. (**B**) Kinetics of

spindle pole separation determined by tubulin immunofluorescence in synchronous meiotic cultures. The same strains as in (A) were analyzed at 25°C and 34°C, and samples were collected at the indicated time points. (**C**) Pulsed-field gel electrophoresis and Southern blot analysis of chromosome III in synchronous meiotic time course of wild type [H7797] and *ycg1-2* mutants [H11281] at 34°C. Samples were collected at the indicated time points. (**D**) Representative images of Smc4-GFP (green) distribution on surface-spread nuclei from wild type [H11471], and *ycg1-2* [H11459] and *smc2-8* mutants [H11455] at 34°C. Chromosomes were counterstained for DNA (DAPI) and Zip1 (red) to mark synapsis. Bar = 1 µm.

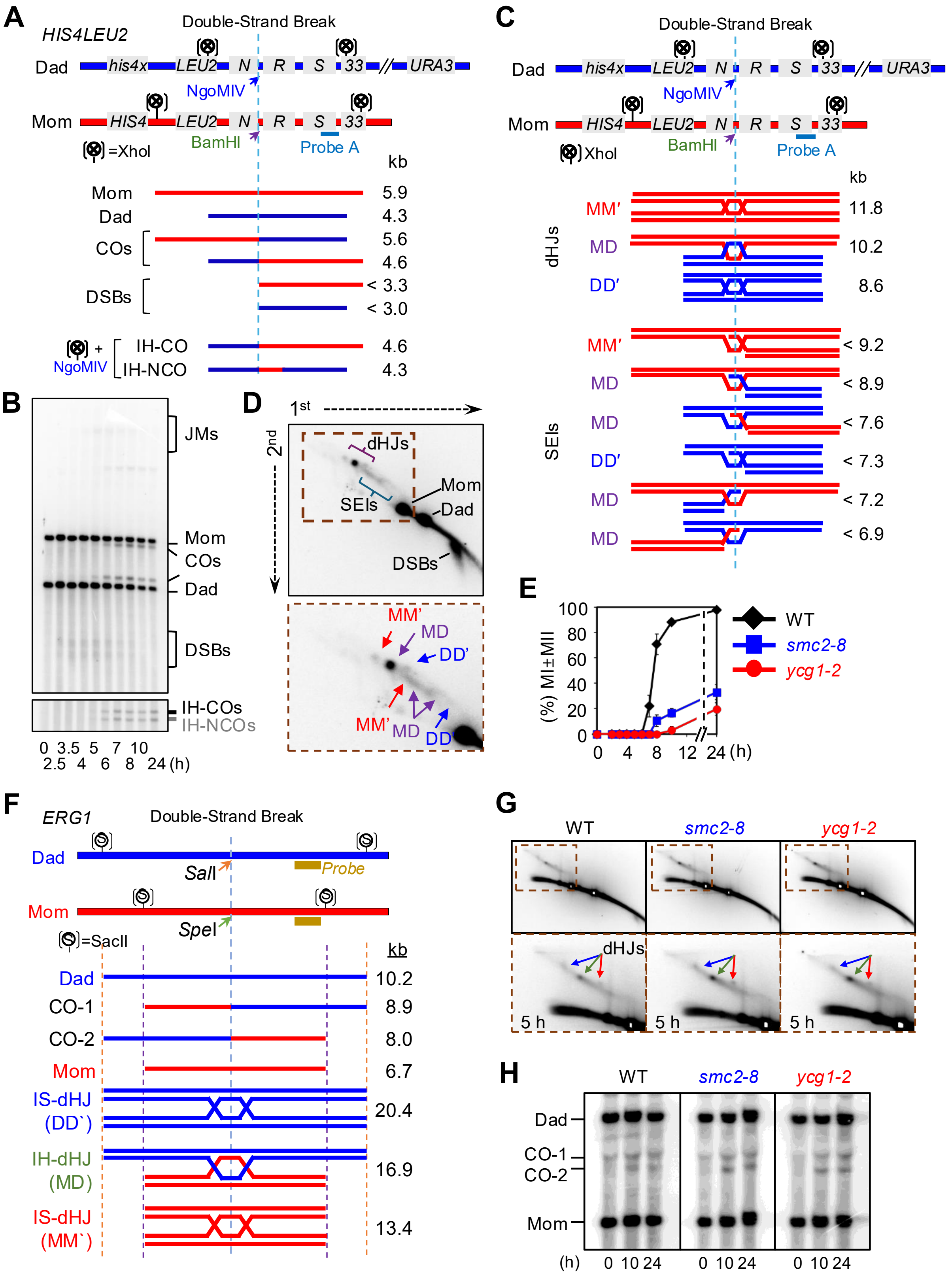
A
HIS4LEU2
Double-Strand Break
Dad
his4x
LEU2
N
R
S
33
URA3
NgoMIV
Mom
HIS4
BamHI
Probe A
=XhoI
kb
Mom 5.9
Dad 4.3
COs 5.6
4.6
DSBs < 3.3
< 3.0
+ NgoMIV
IH-CO 4.6
IH-NCO 4.3
B
JMs
Mom
COs
Dad
DSBs
IH-COs
IH-NCOs
0 3.5 5 7 10
2.5 4 6 8 24 (h)
C
Double-Strand Break
Dad
Mom
NgoMIV
BamHI
Probe A
XhoI
kb
dHJs
MM′ 11.8
MD 10.2
DD′ 8.6
SEIs
MM′ < 9.2
MD < 8.9
MD < 7.6
DD′ < 7.3
MD < 7.2
MD < 6.9
D
1st
2nd
dHJs
SEIs
Mom
Dad
DSBs
MM′
MD
DD′
E
(%) MI±MII
100
80
60
40
20
0
0 4 8 12 24 (h)
WT
smc2-8
ycg1-2
F
ERG1
Double-Strand Break
Dad
SalI
Probe
Mom
SpeI
=SacII
kb
Dad 10.2
CO-1 8.9
CO-2 8.0
Mom 6.7
IS-dHJ (DD`) 20.4
IH-dHJ (MD) 16.9
IS-dHJ (MM`) 13.4
G
WT
smc2-8
ycg1-2
dHJs
5 h
H
WT
smc2-8
ycg1-2
Dad
CO-1
CO-2
Mom
(h) 0 10 24 0 10 24 0 10 24

**Figure S4. Experimental setup for analyzing recombination intermediates at *HIS4LEU2* and *ERG1.* Related to Figure 4.**

(**A**) Schematic of the *HIS4LEU2* locus on chromosome III in Maternal (mom) and paternal (dad) chromosomes, showing XhoI restriction sites and location of probe A. Expected fragment sizes: Mom (5.9 kb), Dad (4.3 kb), crossovers (COs; 5.6 and 4.6 kb), Double-strand breaks (DSBs; < 3.3 and < 3.0 kb), inter-homolog crossover (IH-CO; 4.6kb), and inter-homolog noncrossover (IH-NCO; 4.3 kb). (**B**) Representative 1D gel of wild-type (WT) samples hybridized with probe Across a meiotic time course (0, 2.5, 3.5, 4, 5, 6, 7, 8, 10, and 24 hours) at 34°C. (**C**) Diagrams of recombination intermediates; single-end invasions (SEIs) and double Holliday Junctions (dHJs). Fragment sizes for dHJ species: MM'-dHJs (11.8 kb), MD-dHJs (10.2 kb), DD'-dHJs (8.6 kb). Fragment sizes for SEI species: MM'-SEIs (< 9.0 kb), MD-SEIs (< 8.9, < 7.6, < 7.2, < 6.9 kb), DD'-SEIs (< 7.3 kb). (**D**) Representative 2D gel images of WT showing separation of recombination intermediates by size (first dimension) and shape (second dimension). Red arrows indicate MM'-dHJs and SEIs; purple arrows, indicate MD-dHJs and SEIs, and blue arrows indicate DD'-dHJs and SEIs. (**E**) Analysis of nuclear division profiles of WT [KKY4278], and *smc2-8* [KKY6547] and *ycg1-2* [KKY6544] strains during meiotic time course at 34°C, as determined by DAPI staining. (**F**) Schematic of the *ERG1* locus on chromosome VII for both mom and dad, including the SacII restriction sites and probe location. Expected fragment sizes of parental and recombinant species, as well as dHJs between sisters (MM’, DD’) and homologs (MD), are indicated. (**G**) Representative 2D gel images of WT and condensin mutants showing meiotic recombination intermediates at the *ERG1* locus during meiosis at 34°C. Red, purple, and blue arrows indicate MM'-dHJs, MD-dHJs, and DD'-dHJs, respectively. (**H**) Crossover levels at the *ERG1* locus in WT and condensin mutants at 0, 10, and 24 hours, as determined by 1D gel analysis.

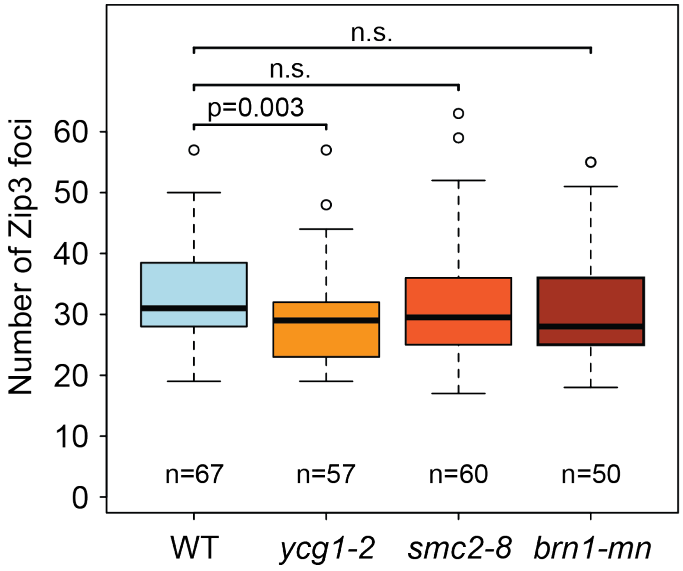


**Figure S5. The number of Zip3-GFP foci is largely unchanged in condensin mutants. Related to Figure 5.**

Analysis of Zip3-GFP foci at pachynema in wild-type [H13116], *ycg1-2* [H13117], *smc2-8* [H13118], and *brn1-mn* [H13141] strains at 34°C. Quantification of the number Zip3-GFP foci per nucleus in the indicated strains. Statistical comparisons were performed using the Wilcoxon rank-sum test with continuity correction.

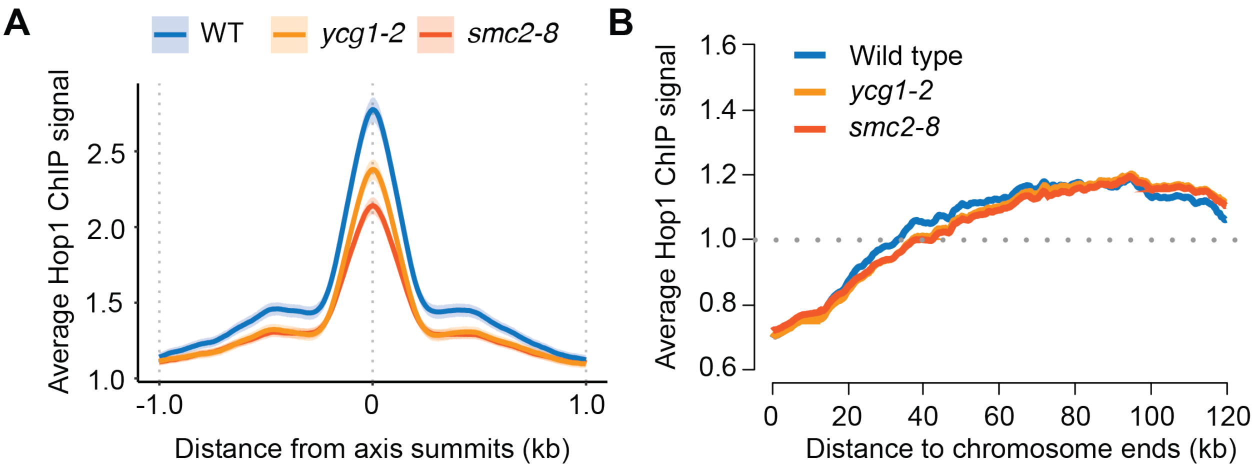


**Figure S6. Mild effects on overall Hop1 enrichment in condensin mutants. Related to Figure 6.**

ChIP-seq analysis of Hop1 binding in wild-type [H7797], *ycg1-2* [H11281], and *smc2-8* [H11364] strains at 34°C, 3 hours after meiotic induction. (**A**) Average Hop1 enrichment at axis attachment sites, as previously defined [S1]. (**B**) Average Hop1 enrichment as a function of distance from chromosome ends. Hop1 overenrichment in the end-adjacent regions (EARs, 20-120kb from telomeres [S2]) is modestly shifted toward more internal chromosomal regions in condensin mutants.

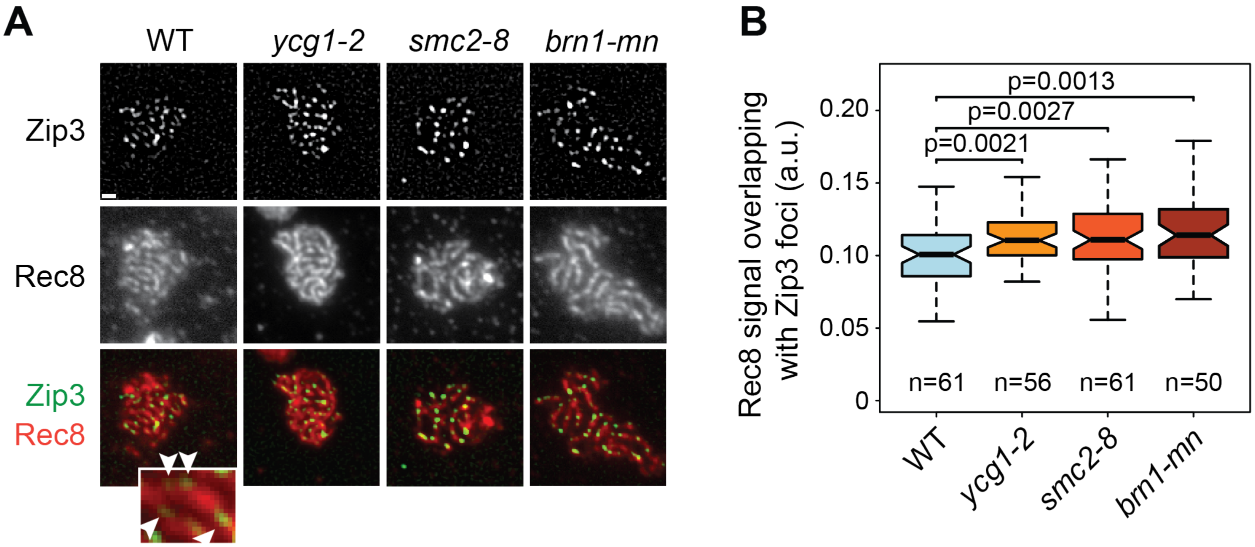


**Figure S7. Increased Rec8 signal at sites of crossover designation. Related to Figure 7.** (**A**) Immunofluorescence analysis of Zip3-GFP (green) and Rec8 (red) on surface-spread pachytene nuclei from wild type [H13116], as well asl *ycg1-2* [H13117], *smc2-8* [H13118], and *brn1-mn* [H13141] mutants, all incubated at 34°C. Arrowheads in zoomed-in detail indicate examples of Zip3 foci that coincide with reduced Rec8 staining in wild type. (**B**) Quantification of Rec8 fluorescence signal colocalizing with Zip3 foci in the indicated genotypes. The number of nuclei analyzed per genotype is indicated. Statistical significance was assessed using Wilcoxon rank-sum test with continuity correction.

**Table S1. Strain list. Related to all figures.**

| Strain Name | Genotype |
|---|---|
| H119 | *MATa/MATα, ho::LYS2/ho::LYS2, lys2/lys2, ura3/ura3, leu2::hisG/ leu2::hisG, his4B::LEU2/his4X::LEU2(Bam)-URA3, arg4-BglII/arg4-Nsp* |
| H4471 | *MATa/MATα, ho::LYS2/ ho::LYS2, lys2/lys2, ura3/ura3, leu2::hisG/ leu2::hisG, his4B::LEU2/his4B::LEU2, arg4-BglII/arg4-BglII, REC8-HA::URA3/REC8-HA::URA3* |
| H6408 | Same as H7797 except *SMC4-PK9::HIS3/SMC4-PK9::HIS3* |
| H7660 | Same as H7797 except *SMC4-PK9::HIS3/SMC4-PK9::HIS3, rec8Δ::HIS3MX6/rec8Δ::HIS3MX6* |
| H7797 | *MATa/MATα, ho::LYS2/ho::LYS2, lys2/lys2, ura3/URA3, leu2::hisG/LEU2, his3::hisG/HIS3, trp1::hisG/TRP1* |
| H8428 | Same as H7797 except *rec8Δ::HIS3MX6/rec8Δ::HIS3MX6* |
| H8601 | *MATa/MATα, ho::LYS2/ ho::LYS2, lys2/lys2, ura3/ura3, leu2::hisG/leu2::hisG, his4X::LEU2-URA3/his4X::LEU2-URA3, ycs4-2::HIS3/ycs4-2::HIS3* |
| H8630 | Same as H7797 except *SMC4-PK9::HIS3/SMC4-PK9::HIS3, spo11-Y135F-HA::URA3/spo11-Y135F-HA::URA3* |
| H8643 | Same as H7797 except *spo11-Y135F-HA::URA3/spo11-Y135F-HA::URA3* |
| H9077 | *MATα/MATa, ho::LYS2/ ho::LYS2, lys2/lys2, HIS3/his3::hisG, URA3/ura3, trp1::hisG/trp1::hisG, leu2::hisG/LEU2, YCS4-13MYC::KanMX/YCS4-13MYC::KanMX* |
| H9441 | Same as H7797 except *SMC4-GFP::TRP1/SMC4-GFP::TRP1, spo11-Y135F-HA::URA3/spo11-Y135F-HA::URA3* |
| H9442 | Same as H7797 except *SMC4-GFP::TRP1/SMC4-GFP::TRP1, rec8Δ::HIS3MX6/rec8Δ::HIS3MX6* |
| H9443 | Same as H7797 except *SMC4-GFP::TRP1/SMC4-GFP::TRP1* |
| H11014 | *MATα/MATa, ho::LYS2/ ho::LYS2, lys2/lys2, HIS3/his3::hisG, URA3/ura3, trp1::hisG/trp1::hisG, leu2::hisG/LEU2, YCS4-13MYC::KanMX/YCS4-13MYC::KanMX, sml1Δ::KanMX/sml1Δ::KanMX, mec1-1/mec1-1* |
| H11058 | *MATα/MATa, ho::LYS2/ ho::LYS2, lys2/lys2, HIS3/his3::hisG, URA3/ura3, trp1::hisG/trp1::hisG, leu2::hisG/LEU2, YCS4-13MYC::KanMX/YCS4-13MYC::KanMX, spo11Δ::TRP1/spo11Δ::TRP1* |
| H11281 | *MATα/MATa, ho::LYS2/ ho::LYS2, lys2/lys2, HIS3/his3::hisG, URA3/ura3, TRP1/trp1::hisG, leu2::hisG/LEU2, ycg1-2::KanMX/ycg1-2::KanMX* |
| H11363 | *MATa/MATα, ho::LYS2/ ho::LYS2, lys2/lys2, ura3/URA3, LEU2/leu2::hisG, HIS3/his3::hisG, TRP/ trp1::hisG, smc4-1/smc4-1* |

| | |
|---|---|
| H11364 | *MATa/ MATα, ho::LYS2/ ho::LYS2, lys2/lys2, URA3/ura3, LEU2/leu2::hisG, HIS3/ his3::hisG, trp1::hisG/TRP1, smc2-8/smc2-8* |
| H11455 | *MATα/MATa, ho::LYS2/ho::LYS2, lys2/lys2, leu2::hisG/LEU2, URA3/ura3, trp1::hisG/trp1::hisG, HIS3/his3::hisG, SMC4-GFP::TRP1/SMC4-GFP::TRP1, smc2-8/smc2-8* |
| H11459 | *MATα/MATa, ho::LYS2/ho::LYS2, lys2/lys2, leu2::hisG/LEU2, URA3/ura3, trp1::hisG/trp1::hisG, HIS3/his3::hisG, SMC4-GFP::TRP1/SMC4-GFP::TRP1, ycg1-2::KanMX /ycg1-2::KanMX* |
| H11471 | *MATα/MATa, ho::LYS2/ho::LYS2, lys2/lys2, leu2::hisG/LEU2, URA3/ura3, trp1::hisG/ trp1::hisG, HIS3/his3::hisG, SMC4-GFP::TRP1/SMC4-GFP::TRP1* |
| H11499 | *MATa/MATα, ho::LYS2/ho::LYS2, lys2/lys2, trp1::hisG/trp1::hisG, his3::hisG/HIS3, LEU2/leu2::hisG, ura3/URA3, zip1Δ::LYS2/zip1Δ::LYS2, SMC4-GFP::TRP1/SMC4-GFP::TRP1* |
| H11735 | *MATa/MATα, ho::LYS2/ho::LYS2, lys2/lys2, LEU2/leu2::hisG, HIS3/his3::hisG, trp1::hisG/trp1::hisG, ura3/URA3, SMC4-GFP::TRP1/SMC4-GFP::TRP1, MSH4-13myc::KanMX6/MSH4-13myc::KanMX6* |
| H11758 | *MATα/MATa, ho::LYS2/ho::LYS2, lys2/lys2, URA3/ura3, LEU2/leu2::hisG, HIS3/his3::hisG, TRP1/trp1::hisG, pch2Δ::KanMX/pch2Δ::KanMX* |
| H12022 | *MATa/MATα, ho::LYS2/ho::LYS2, lys2/lys2, URA3/ura3, leu2::hisG/LEU2, HIS3/his3::hisG, trp1::hisG/TRP1, pCLB2-HA-BRN1::KanMX4/pCLB2-HA-BRN1::KanMX4* |
| H12137 | *MATa/MATα, ho::LYS2/ho::LYS2, lys2/lys2, ura3/URA3, his3::hisG/HIS3, trp1::hisG/TRP1, LEU2/leu2::hisG, mek1Δ::kanMX/mek1Δ::kanMX* |
| H12173 | *MATα/MATa, ho::LYS2/ho::LYS2, lys2/lys2, ura3/ura3, LEU2/leu2::hisG, his3::hisG/HIS3, trp1::hisG/TRP1, spo11-Y135F-HA-URA3/spo11-Y135F-HA-URA3* |
| H12194 | *MATα/MATa, ho::LYS2/ho::LYS2, lys2/lys2, URA3/ura3, HIS3/his3::hisG, trp1::hisG/TRP1, leu2::hisG/LEU2, mek1Δ::kanMX/mek1Δ::kanMX, ycg1-2::KanMX/ycg1-2::KanMX* |
| H12205 | *MATa/MATα, ho::LYS2/ho::LYS2, lys2/lys2, URA3/ura3, trp1::hisG/TRP1, HIS3/his3::hisG, LEU2/leu2::hisG, mek1Δ::kanMX/mek1Δ::kanMX, smc2-8/smc2-8* |
| H12227 | *MATa/MATα, ho::LYS2/ho::LYS2, lys2/lys2, ura3/ura3, trp1::hisG/TRP1, his3::hisG/his3::hisG, LEU2/leu2::hisG, ycg1-2::KanMX/ ycg1-2::KanMX, spo11-Y135F-HA-URA3/spo11-Y135F-HA-URA3* |
| H12245 | *MATa/MATα, ho::LYS2/ho::LYS2, lys2/lys2, ura3/ura3, LEU2/LEU2, his3::hisG/HIS3, trp1::hisG/trp1::hisG, smc2-8/smc2-8, spo11-Y135F-HA-URA3/spo11-Y135F-HA-URA3* |

| | |
|---|---|
| H12818 | *MATα/MATa, ho::LYS2/ho::LYS2, lys2/lys2, URA3/ura3, his3::hisG/his3::hisG, leu2::hisG/LEU2, trp1::hisG/trp1::hisG, ecm11Δ::HIS3/ecm11Δ::HIS3, SMC4-GFP::TRP1/SMC4-GFP::TRP1* |
| H12925 | *MATα/MATa, ho::LYS2/ho::LYS2, lys2/lys2, ura3/URA3, leu2::hisG/LEU2, his3::hisG/HIS3, trp1::hisG/trp1::hisG, zip3Δ::KanMX6/zip3Δ::KanMX6, SMC4-GFP::TRP/SMC4-GFP::TRP1* |
| H12926 | *MATα/MATa, ho::LYS2/ho::LYS2, lys2/lys2, ura3/URA3, LEU2/leu2::hisG, his3::hisG/HIS3, trp1::hisG/trp1::hisG, ZIP3-13myc::hphMX4/ZIP3-13myc::hphMX4, SMC4-GFP::TRP1/SMC4-GFP::TRP1* |
| H13116 | *MATα/MATa, ho::LYS2/ho::LYS2, lys2/lys2, LEU2/leu2::hisG, HIS3/his3::hisG, TRP1/trp1::hisG, ura3/ura3, ZIP3-GFP::URA3/+* |
| H13117 | *MATa/MATα, ho::LYS2/ho::LYS2, lys2/lys2, ura3/ura3, TRP1/trp1::hisG, his3::hisG/HIS3, LEU2/LEU2, ycg1-2::KanMX/ycg1-2::KanMX, ZIP3-GFP::URA3/+* |
| H13118 | *MATα/MATa, ho::LYS2/ho::LYS2, lys2/lys2, LEU2/leu2::hisG, HIS3/his3::hisG, TRP1/trp1::hisG, ura3/ura3, smc2-8/smc2-8, ZIP3-GFP::URA3/+* |
| H13141 | *MATα/MATa, ho::LYS2/ho::LYS2, lys2/lys2, LEU2/leu2::hisG, HIS3/his3::hisG, trp1::hisG/TRP1, ura3/ura3, pCLB2-HA-BRN1::KanMX4/pCLB2-HA-BRN1::KanMX4, ZIP3-GFP::URA3/+* |
| H13167 | *MATα/MATa, ho::LYS2/ho::LYS2, lys2/lys2, leu2::hisG/LEU2, URA3/ura3, trp1::hisG/ trp1::hisG, his3::hisG/his3::hisG, SMC4-GFP::TRP1/SMC4-GFP::TRP1, zip2Δ::HIS3/zip2Δ::HIS3* |
| H13168 | *MATα/MATa, ho::LYS2/ho::LYS2, lys2/lys2, leu2::hisG/LEU2, ura3/ura3, trp1::hisG/ trp1::hisG, his3::hisG/HIS3, SMC4-GFP::TRP1/SMC4-GFP::TRP1, msh4Δ::URA3/msh4Δ::URA3* |
| H13208 | *MATα/MATa, ho::LYS2/ ho::LYS2, lys2/lys2, HIS3/HIS3, URA3/ura3, trp1::hisG/TRP1, leu2::hisG/LEU2, YCS4-13MYC::KanMX/YCS4-13MYC::KanMX, sml1Δ::KanMX/sml1Δ::KanMX* |
| H13306 | *MATα/MATa, ho::LYS2/ho::LYS2, lys2/lys2, leu2::hisG/LEU2, URA3/ura3, trp1::hisG/ trp1::hisG, his3::hisG/his3::hisG , SMC4-GFP::TRP1/SMC4-GFP::TRP1, msh5Δ::HIS3/msh5Δ::HIS3* |
| H13307 | *MATα/MATa, ho::LYS2/ho::LYS2, lys2/lys2, leu2::hisG/LEU2, URA3/ura3, trp1::hisG/ trp1::hisG, his3::hisG/his3::hisG , SMC4-GFP::TRP1/SMC4-GFP::TRP1, zip4Δ::HIS3/zip4Δ::HIS3* |
| H13373 | *MATα/MATa, ho::LYS2/ ho::LYS2, lys2/lys2, HIS3/his3::hisG, URA3/ura3, trp1::hisG/TRP1, leu2::hisG/LEU2, YCS4-13MYC::KanMX/YCS4-13MYC::KanMX, zip1Δ::LYS2/zip1Δ::LYS2* |
| H13376 | *MATα/MATa, ho::LYS2/ ho::LYS2, lys2/lys2, his3::hisG/his3::hisG, URA3/ura3, trp1::hisG/TRP1, leu2::hisG/LEU2, YCS4-13MYC::KanMX/YCS4-13MYC::KanMX, zip2Δ::HIS3/zip2Δ::HIS3* |

| | |
|---|---|
| H13379 | *MATα/MATa, ho::LYS2/ ho::LYS2, lys2/lys2, his3::hisG/his3::hisG, URA3/ura3, trp1::hisG/TRP1, leu2::hisG/LEU2, YCS4-13MYC::KanMX/YCS4-13MYC::KanMX, zip4Δ::HIS3/zip4Δ::HIS3* |
| H13382 | *MATα/MATa, ho::LYS2/ ho::LYS2, lys2/lys2, HIS3/his3::hisG, ura3/ura3, TRP1/TRP1, LEU2/LEU2, YCS4-13MYC::KanMX/YCS4-13MYC::KanMX, msh4Δ::URA3/msh4Δ::URA3* |
| H13391 | *MATα/MATa, ho::LYS2/ ho::LYS2, lys2/lys2, his3::hisG/HIS3, URA3/ura3, trp1::hisG/TRP1, leu2::hisG/LEU2, YCS4-13MYC::KanMX/YCS4-13MYC::KanMX,* |
| H13394 | *MATα/MATa, ho::LYS2/ ho::LYS2, lys2/lys2, HIS3/his3::hisG, URA3/ura3, trp1::hisG/TRP1, leu2::hisG/LEU2, YCS4-13MYC::KanMX/YCS4-13MYC::KanMX, zip3Δ::KanMX/zip3Δ::KanMX* |
| KKY2945 | *MATα/MATa, ho::hisG/ho::hisG, leu2::hisG/leu2::hisG, ura3/ura3, HIS4::LEU2/his4-X::LEU2-URA3, ERG1::SalI/ERG1::SpeI* |
| KKY4278 | *MATα/MATa, ho::hisG/ho::hisG, leu2::hisG/leu2::hisG, ura3/ura3, HIS4::LEU2/his4-X::LEU2-URA3* |
| KKY6547 | *MATα/MATa, ho::hisG/ho::hisG, leu2::hisG/leu2::hisG, ura3/ura3, HIS4::LEU2/his4-X::LEU2-URA3, smc2-8/smc2-8* |
| KKY6544 | *MATα/MATa, ho::hisG/ho::hisG, leu2::hisG/leu2::hisG, ura3/ura3, HIS4::LEU2/his4-X::LEU2-URA3, ycg1-2::KanMX/ycg1-2::KanMX* |
| KKY6850 | *MATα/MATa, ho::hisG/ho::hisG, leu2::hisG/leu2::hisG, ura3/ura3, HIS4::LEU2/his4-X::LEU2-URA3, ERG1::SalI/ERG1::SpeI, smc2-8/smc2-8* |
| KKY6851 | *MATα/MATa, ho::hisG/ho::hisG, leu2::hisG/leu2::hisG, ura3/ura3, HIS4::LEU2/his4-X::LEU2-URA3, ERG1::SalI/ERG1::SpeI, ycg1-2::KanMX/ycg1-2::KanMX* |
