## Supplementary material for "Crossover designation recruits condensin to reorganize the meiotic chromosome axis": Key resources table

| REAGENT or RESOURCE | SOURCE | IDENTIFIER |
|---|---|---|
| Antibodies | | |
| Anti-V5 Agarose Affinity Gel antibody | Sigma | Cat#A7345; RRID: AB_10062721 |
| Rat monoclonal anti-HA (3F10 ) | Roche Applied Science | Cat#11867431001; RRID: AB_390919 |
| Rabbit polyclonal anti-Red1 | Kind gift from N. Hollingsworth [102] | N/A |
| Rabbit polyclonal anti-Hop1 | Kind gift from N. Hollingsworth [103] | N/A |
| Guinea Pig polyclonal anti-Rec8 | Kind gift from N. Hollingsworth [107] | N/A |
| Goat polyclonal anti-Zip1 (yC-19) | Santa Cruz Biotechnology | Cat#sc-15632; RRID: AB_672952 |
| Chicken polyclonal anti-GFP | Abcam | Cat#13970; RRID: AB_300798 |
| Mouse monoclonal anti-Nop1 | EnCor Technologies | Cat#MCA-28F2; RRID: AB_1077286 |
| Rabbit polyclonal anti-Pch2 | Kind gift from A. Shinohara [7] | N/A |
| Rabbit polyclonal anti-Rec8-pS521 | Kind gift from G. Brar [108] | N/A |
| Rat monoclonal anti-Tubulin YOL1/34 | Santa Cruz Biotechnology | Cat#sc-53030; RRID: AB_2272440 |
| Mouse anti-Myc 4A6 | Millipore | Cat#05724; RRID: AB_4169237 |
| Rabbit anti-Myc 4A6 | Cell Signaling Technology | Cat#2272S; RRID: AB_10692100; lot6 |
| Rabbit anti-Fpr3 | Kind gift from J. Thorner [109] | N/A |
| Secondary anti-Rabbit Alexa 488 | Jackson Immunoresearch | Cat#711-545-152; RRID: AB_139808 |
| Secondary anti-Rabbit Cy3 | Jackson Immunoresearch | Cat#711-165-152; RRID: AB_125364 |
| Secondary anti-Rabbit Cy5 | Jackson Immunoresearch | Cat#711-175-152; RRID:AB_150312 |
| Secondary anti-Chicken Alexa 488 | Jackson Immunoresearch | Cat#703-545-155; RRID: AB_2340375 |
| Secondary anti-Mouse Alexa 488 | Jackson Immunoresearch | Cat#715-545-150; RRID: AB_173495 |
| Secondary anti-Mouse Cy5 | Jackson Immunoresearch | Cat#715-175-151; RRID: AB_165683 |
| Secondary anti-Rat FITC | Jackson Immunoresearch | Cat#712-095-153; RRID: AB_170320 |
| Secondary anti-Goat Alexa 488 | Jackson Immunoresearch | Cat#705-545-003; RRID: AB_174093 |
| Secondary anti-Goat Cy3 | Jackson Immunoresearch | Cat#705-166-147; RRID: AB_168948 |
| Secondary anti-Guinea Pig 488 | Jackson Immunoresearch | Cat#706-545-148; RRID: AB_160322 |

| | | |
|---|---|---|
| Secondary anti-Guinea Pig Cy3 | Jackson Immunoresearch | Cat#706-165-148; RRID: AB_161782 |
| Secondary anti-Guinea Pig Cy5 | Jackson Immunoresearch | Cat#706-175-148; RRID: AB_170461 |
| Digital anti-Mouse HRP | Kindle Biosciences | Cat#R1005; RRID: AB_2800463 |
| Vibrance antifade Mounting Medium with DAPI | Vectashield | H-1800-10 |
| Chemicals, peptides, and recombinant proteins | | |
| Zymolase T100 | US Biological | Z1004 |
| RNaseA | Sigma Aldrich | S4642 |
| Formaldehyde solution | Sigma Aldrich | F1635 |
| $\alpha$-$^{32}$P-dCTP | REVVITY | BLU513H250UC |
| Formaldehyde granular | VWR | 100504-160 |
| Glycine | Sigma-Aldrich | G7126 |
| PMSF | Sigma Aldrich | P7626 |
| GE Gammabind Sepharose | GE Healthcare | 17-0885-01 |
| Glycogen | Ambion | AM9510 |
| Proteinase K | Roche | 3115828001 |
| dNTP Mixture (each 2.5 mM) | Takara | 4030 |
| T4 Polynucleotide Kinase (10,000 units/ml) | New England BioLabs | M0201S |
| Klenow Fragment | New England BioLabs | M0210S |
| T4 DNA polymerase | New England BioLabs | M0203S |
| dATP (100mM) | ThermoFisher Scientific | 10216018 |
| Klenow Fragment exo- | New England BioLabs | M0212S |
| Agencourt AMPure XP | Beckman Coulter | A63880 |
| Quick Ligase | New England BioLabs | M2200S |
| dNTP Mix (each 10mM) | ThermoFisher Scientific | R0192 |
| Phusion High-Fidelity DNA Polymerase | ThermoFischer Scientific | F-530XL |
| SeaKem Agarose | Lonza (via Fisher Scientific) | BMA50004 |
| TrackIt 100bp DNA ladder | ThermoFisher Scientific | 10488-058 |
| Antifoam 204 | Sigma-Aldrich | A8311 |
| Bacto-agar | BD Biosciences | 214010 |
| Bacto-peptone | Gibco | 211677 |
| Bacto-yeast extract | Gibco | 212750 |
| D-sorbitol | Duksan | 849 |
| dCTP ($\alpha$-$^{32}$P) | Revvity | PK-BLU013H |
| Dextrose | BD Biosciences | 215530 |
| D-(+)-Raffinose pentahydrate | MP Biomedicals | 102797 |
| EDTA disodium dihydrate | Duchefa Biochemie | E0511 |
| Ethidium bromide | Sigma-Aldrich | E8751 |
| Guanidine-HCl | Duksan | BG003-1 |
| *Xho*I | Enzynomics | R007 |

| | | |
|---|---|---|
| *Sac*II | Enzynomics | R008 |
| Lipsol | Scilabware | Py.40023 |
| Lyticase | Sigma-Aldrich | L4025 |
| *Ngo*MIV | Enzynomics | R095 |
| Phenol/chloroform/isoamyl alcohol (PCI) | Bio-Solution | BP026 |
| Potassium hydrogen phthalate | Daejung | 6581-4400 |
| Proteinase K | Enzynomics | PR003 |
| Ribonuclease A | Sigma-Aldrich | R6513 |
| SeaKem Gold Agarose | Lonza | 50152 |
| Sodium acetate | Samchun Chemicals | S0292 |
| Sodium azide | Samchun Chemicals | S0317 |
| Sodium chloride | Duchefa Biochemie | S0520 |
| Sodium dodecyl sulfate (SDS) | Duchefa Biochemie | S1377 |
| Sodium hydroxide (NaOH) | Samchun Chemicals | S0620 |
| Sodium lauroyl sarcosinate | Sigma-Aldrich | L9150 |
| Sucrose | Sigma-Aldrich | S0389 |
| UltraKem LE Agarose | Young Science | Y50004 |
| yeast nitrogen base without amino acids and ammonium sulfate | BD Bioscience | 291940 |
| β-Mercaptoethanol | Bio-Rad | 16107710 |
| Biodyne® B Membrane, 0.45 µm | Pall corporation | 6028 |
| Critical commercial assays | | |
| Qubit dsDNA HS Assay Kit | Life Technologies (Invitrogen) | Q32851 |
| MinElute PCR Purification Kit | Qiagen | 28004 |
| Agilent TapeStation HS D1000 (DNA) Reagents | Agilent | 5067-5585 |
| Agilent TapeStation HS D1000 (DNA) Tape | Agilent | 5067-5584 |
| Qiaquick gel extraction kit (50 columns) | Qiagen | 28704 |
| KAPA Library Quantification Kit - Complete Kit (Universal) | Roche Applied Science | 7960140001 |
| Prime-It RmT Random Primer Labeling Kit | Agilent | 300392 |
| EZ™-Random primer DNA labeling kit | Enzynomics | Cat# EZ022S |
| Deposited data | | |
| Meiotic ChIP-seq data | This study | GSE154722 |
| Asynchronous ChIP-seq data | Paul et al. 2018 [37] | GSE106104 |
| Experimental models: Organisms/strains | | |
| See Table S1 for the strains | N/A | N/A |
| Oligonucleotides | | |
| *MRC1* locus amplification Forward: 5'- ttc gag gag tca att tcc cgt ttc -3'<br>*MRC1* locus amplification Reverse: 5'- cat cag gaa cag aaa acc caa ata g -3' | Blitzblau et al. ,2007 [110] | Primer321, Primer322 |
| *MRC1* nested probe amplification Forward: 5'- gag agt ctt gcc ttc agg -3'<br>*MRC1* nested probe amplification Reverse: 5'- gga tgt acc acc aca gac -3' | Blitzblau et al., 2007 [110] | Primer330, Primer331 |

| | | |
|---|---|---|
| *HIS4LEU2* locus, 5 -ATATACCGGTGTTGGGCCTT T-3 and 5 -ATATAGATCTCCTACAATATCAT-3 | Kim et al., 2010 [100] | |
| *ERG1* locus, 5 -ATGGAAGATATAGAAGGATACGAACC-3 and 5 -GCGACGCAAATTCGCCGATGGTTTG-3 | Lee et al., 2021 [64] | |
| Software and algorithms | | |
| R | R Core Team | https://www.R-project.org/ |
| R studio | Rstudio | https://www.rstudio.com |
| Fiji | ImageJ | https://fiji.sc |
| DiAnA plugin | Gilles et al., 2017 [115] | https://imagej.net/plugins/distance-analysis |
| Bowtie | Langmead et al., 2009 [112] | |
| Bowtie 2 | Langmead et al., 2012 [113] | http://bowtie-bio.sourceforge.net/bowtie2/index.shtml |
| Trimmomatic | | http://www.usadellab.org/cms/index.php?page=trimmomatic |
| MACS v2.1.1 | Zhang et al., 2008 [114] | (https://github.com/taoliu/MACS |
| BD Accuri C6 software | BD Biosciences | C6 Plus Analysis Software for PC or Mac |
| FlowJo™ v10.8 | BD Biosciences | FlowJo™ Software \| BD Biosciences |
| Graphpad prism 10 | Graphpad | https://www.graphpad.com/ |
| Quantity One | Bio-Rad | Quantity One 1-D Analysis Software \| Bio-Rad |
| Other | | |
| BD Accuri™ C6 Plus Flow Cytometer | BD Biosciences | https://www.bdbiosciences.com/ko-kr/products/instruments/flow-cytometers/research-cell-analyzers/bd-accuri-c6-plus |
| BD FACSCalibur™ Flow Cytometer | BD Bioscience | https://www.bdbiosciences.com/en-us/products/instruments/flow-cytometers/clinical-cell-analyzers/calibur-discontinuation |
| Personal molecular imager | Bio-Rad | Cat# 170-9400 |
